# Villin-SNAP25 interaction regulates insulin secretion downstream of ICA512 signaling

**DOI:** 10.64898/2026.01.13.699220

**Authors:** Hassan Mziaut, Preethi Kesavan, Chunguang Chen, Ali Gheisari, Jaber Dehghany, Andreas Műller, Anke Sönmez, Katharina Ganß, Inna Kalaidzidis, Carla Münster, Manon von Bülow, Matthias Lohmann, Yannis Kalaidzidis, Anke M. Schulte, Raphael Scharfmann, Stefan Lehr, Hadi Al-Hasani, Hella Hartmann, Michael Meyer-Hermann, Stephan Speier, Michele Solimena

## Abstract

Pancreatic islet β-cells step-up insulin production and secretion in response to the elevation of blood glucose levels. ICA512/PTPRN is a transmembrane cargo of the insulin secretory granules (SGs). Upon SG exocytosis, its Ca^2+/^calpain mediated cleavage at the plasma membrane generates an ICA512-cleaved cytosolic fragment (ICA512-CCF) which as a decoy phosphatase prolongs phospho-STAT5/3 activities, thereby enhancing the transcription of mRNAs for SG cargoes, including *insulin* and its own, to replenish SG stores. In addition, ICA512 positively regulates the expression of F-actin modifier *villin*. Villin, in turn, modulates the size of actin cages surrounding cortical SGs, hence regulating SG mobility and exocytosis. Here we show that villin controls the number of SG docking sites for exocytosis by directly interacting at low glucose with the t-SNARE SNAP-25, thus restricting in this condition SG access to fusion sites and insulin release. Replacement of ICA512-CCF N-terminal ubiquitin-acceptor lysine 609 (K609) with valine (V) stabilized ICA512-CCF in insulinoma cells and homozygous *ICA512^K609V^* mice and enhanced the levels of phospho-STAT5/3 and villin. In *ICA512^K609V^* female mice these molecular traits correlated with reduced body weight, improved insulin sensitivity, reduced basal insulin secretion and onset time for glucose stimulated insulin secretion. Taken together, these data demonstrates that SG exocytosis induced ICA512 retrograde signaling acts in concert with the novel SNARE complex regulator villin to plastically adapt β-cell actin cytoskeleton and access to SG docking sites for optimal control of insulin secretion in response to variations in extracellular glucose levels.

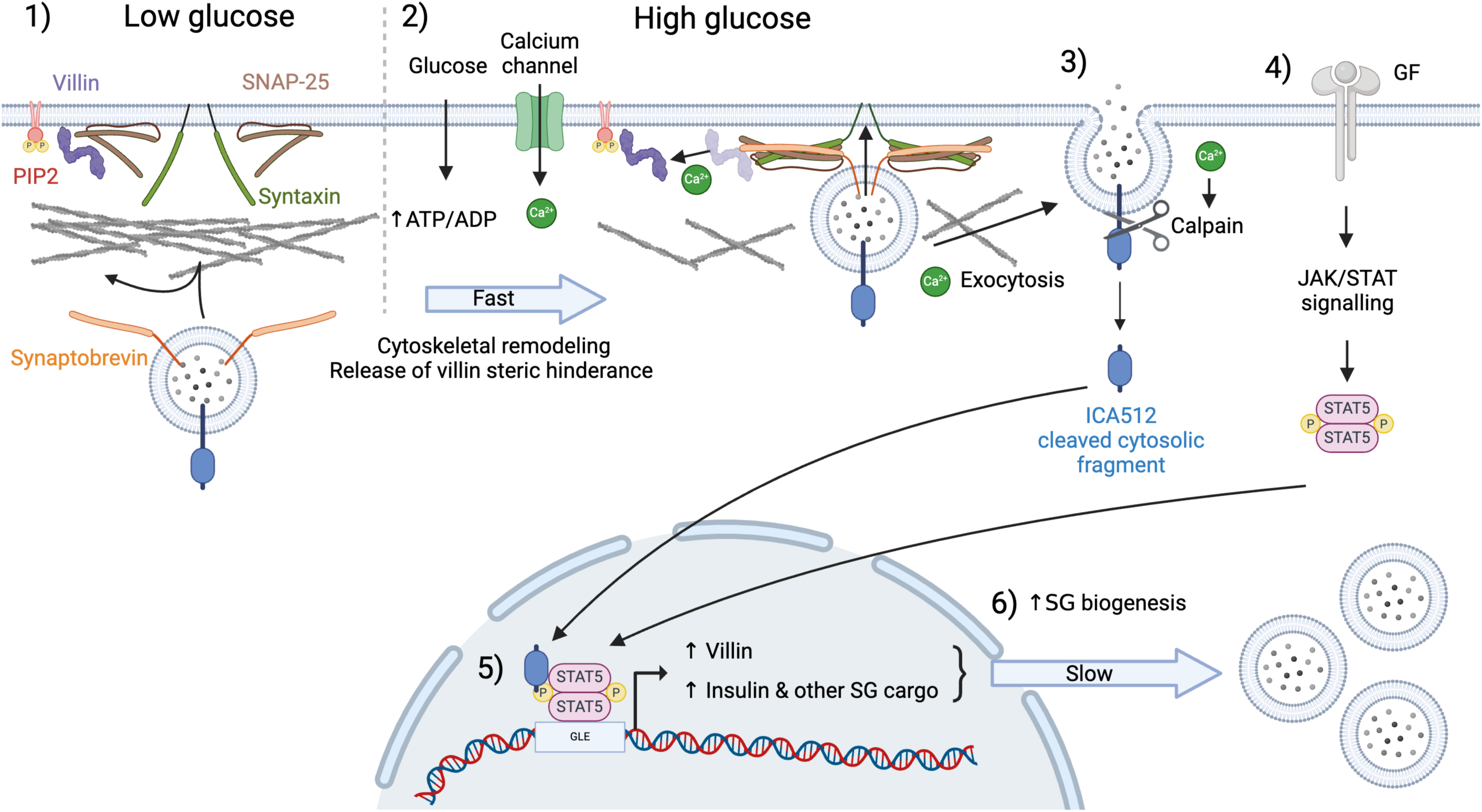
Graphical abstract. Model of ICA512 and STAT3/5–dependent regulation of β-cell function. At basal conditions, villin limits secretory granule (SG) docking by interacting with SNAP-25, restricting insulin release (1). Increased metabolic load triggers hyperglycemia-induced ICA512 processing and growth factor–mediated STAT3/5 phosphorylation (2–4), enhancing ISG mRNAs, granule biogenesis, and villin expression (5,6). Villin modulates SNARE protein availability, controlling SG docking and secretion. Together, ICA512 retrograde signaling and villin-mediated SNARE regulation dynamically remodel the β-cell actin cytoskeleton, adjusting SG docking site access to ensure appropriate insulin secretion in response to fluctuating glucose levels.

## Introduction

Regulated exocytosis enables endocrine cells and neurons to secrete peptide hormones, neuropeptides, and other molecules stored in vesicles called secretory granules (SGs)/large dense core vesicles in response to stimulation. Endocrine β-cells of the pancreatic islets produce and secrete the insulin hormone to lower blood glucose levels upon their elevation and thus restore metabolic homeostasis. Accordingly, the biogenesis and exocytosis of insulin containing SGs must be strictly controlled to prevent blood glucose from reaching dangerously low or high levels, as in the case of diabetes mellitus.

In β-cells, cortical actin microfilaments modulate the peripheral myosin-driven transport of SGs and their exocytosis. Organized as a dense web beneath the plasma membrane (Orci et al, 1972), F-actin depolymerizes upon hyperglycemia to facilitate the access of SGs to exocytic sites and insulin secretion (Thurmond et al, 2003). Disruption of F-actin, using compounds like cytochalasin B (Jijakli et al, 2002), latrunculin B (Thurmond et al, 2003) or Clostridium Botulinum C2 toxin (Li et al, 1994), acutely enhances insulin secretion. Reduced expression of gelsolin, an actin modifier, perturbs F-actin remodeling and reduces glucose-stimulated insulin secretion (GSIS) (Tomas et al, 2006). Similarly, depletion of villin, a gelsolin-related actin modifier, reduces GSIS by enlarging F-actin cages around insulin SGs, thus increasing their motility and basal insulin release (Mziaut et al, 2016).

Villin expression in rodent β-cells is regulated by islet cell autoantigen (ICA512) (Mziaut et al, 2016), a pseudo receptor protein tyrosine phosphatase enriched in SG membranes (Solimena et al, 1996). Upon insulin SG exocytosis, ICA512 transiently translocates to the plasma membrane, where its intracellular tail is cleaved by Ca²⁺/calpain, generating an ICA512-cleaved cytosolic fragment (ICA512-CCF), which includes the catalytically inactive PTP domain. Acting as a decoy phosphatase, ICA512-CCF prevents phospho-STAT5/3 (PY-STAT5/3) dephosphorylation and inactivation (Mziaut at al, 2006; Mziaut et al, 2008). Hence, SG exocytosis-driven generation of ICA512-CCF combined with growth-hormone-induced STAT5/3 phosphorylation, increases the transcription of mRNAs encoding for SG cargoes such as insulin and ICA512, aiding insulin store replenishment. In partially pancreatectomized mice, convergence of ICA512 and STAT5 signalling fostered β-cell regeneration as a complementary mean to restore insulin production (Mziaut et al, 2008). Based on these data, we postulated that synergy of ICA512 and STAT5/3 signaling enables β-cells to adjust insulin supply to metabolic needs. However, most evidence for this model was obtained from analyses conducted in insulinoma cells, as the occurrence of ICA512-CCF and it’s signaling in primary β-cells remained to be shown. Moreover, how ICA512 regulates villin expression also remains unclear.

To close this knowledge gap and gain further mechanistic insight into how villin and ICA512 jointly modulate GSIS we combined in silico modeling with experimental studies in vitro and in vivo. We now show that villin binds to the t-SNARE SNAP25, hence dynamically regulating not only the mobility of insulin SGs, but also their docking to sites for exocytosis. villin and ICA512 expression increase in parallel upon induced differentiation of human EndoC-βH3 β-like cells, conceivably to tune the plastic remodeling of the cortical actin cytoskeleton in accordance to the expansion of the SG stores. Exploitation of a knock-in mouse in which single point mutagenesis of ICA512 enabled the stabilization of short-lived ICA512-CCF provides conclusive evidence for the ability of ICA512 retrograde signaling to positively regulate villin expression, insulin secretion and metabolic homeostasis.

## Results

### *Villin* depletion increases the number of docking sites accessible to SGs

We showed that depletion of villin in insulinoma cell lines and primary β-cells blunts GSIS due to increased basal insulin release (Mziaut et al, 2016). Based on TIRF imaging analyses, we attributed this phenotype to the greater mobility of insulin SGs in *villin*-depleted (*Vil*-siRNA) insulinoma cells exposed to resting glucose levels (2.8 mM) compared to control cells. This increased mobility, in turn, was explained with the larger size of the actin cages restricting cortical SGs due to the reduced villin F-actin bundling activity. To investigate further the relationship between SG mobility and regulated exocytosis, we developed an *in-silico* model (Dehghany et al, 2015), in which multiple parameters were adjusted based on experimental data about insulin SG biosynthesis, distribution and dynamics in resting vs. glucose stimulated β-cells (Rorsman and Renström, 2003; Baltrusch et al, 2008; Fava et al, 2012). We have previously shown that the measured second-phase insulin secretion could only be reproduced by the mathematical model when the number of docking sites (DSs) was dynamic (Dehghany et al, 2015). In the original model (Dehghany et al, 2015), the number of DSs was assumed to be directly and solely regulated by the glucose concentration. However, the dynamics of SGs pertained measurements about their motility on microtubules and actin filaments as well as their rate of docking and fusion at the plasma membrane which in turn include the involvement of SNARE and other accessory proteins. Here we applied this algorithm to separately examine the contribution of various parameters to the observed behavior, i.e. blunted GSIS upon villin depletion (Mziaut et al, 2016). However, since villin affects the global shape of F-actin, which in turn can affect SG biogenesis and dynamics, the model was updated to also consider other parameters, such as the number of accessible docking sites for SG exocytosis. Implementing a variable number of available docking sites resulted in a model predicting that villin depletion alters the cortical cytoskeleton architecture such that the number of SG docking in resting β-cells is enhanced (Fig. 1a-b). Based on this prediction, we investigated the potential relationship of villin and SNARE proteins for regulated exocytosis. In *vil-*siRNA INS-1 cells exposed to 2.8 mM glucose the number of insulin SGs visualized with the TMR-labeled Insulin-SNAP reporter (INS-SNAP^TMR^) (Ivanova et al, 2013) which colocalized with Syntaxin1-GFP (Stx1-GFP) as a SNARE marker of docking sites was increased (Fig. 1c-d). This increase was reversed by expression of mouse villin with a construct resistant to *vil*-siRNA (Fig. 1e and Extended Data Fig. 1). Analysis with Motion tracking, as described (Hoboth et al, 2015), showed that the percentage of nearly immobile INS-SNAP^TMR+^ SGs colocalized with Stx1-GFP^+^ decreased overtime, conceivably due to exocytosis, but remained higher in *vil*-siRNA INS-1 cells (Fig. 1f). This data corroborates the hypothesis that villin depletion increases access of SGs to docking sites.

**Figure 1.**
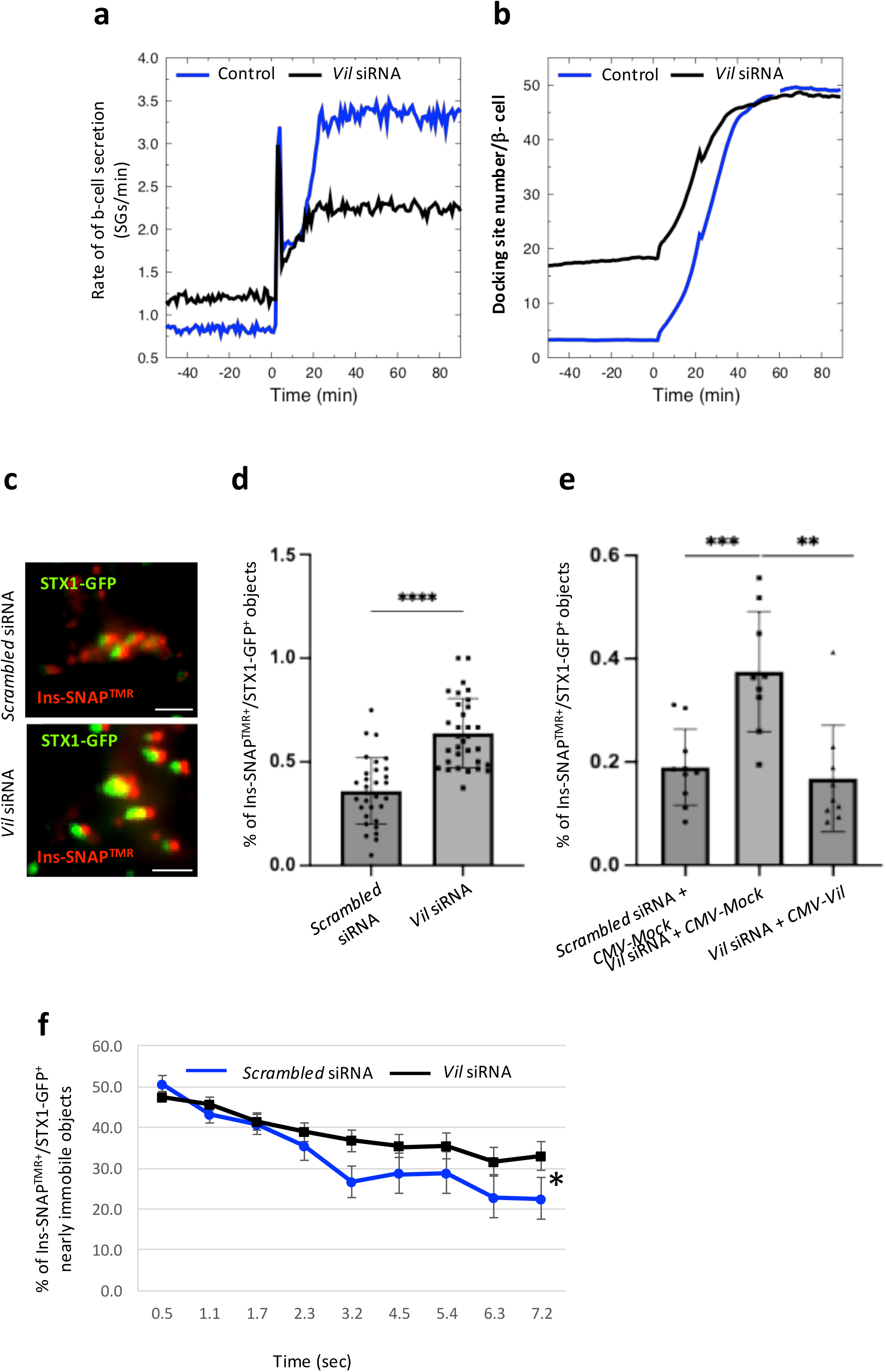
Depletion of villin increases docking sites number. (a-b) In *silico* model of GSIS in relationship to the insulin secretion rate (a) and the number of docking sites (b) in INS-1 cells treated with scrambled or *villin (Vil)* siRNAs in response to an increase in extracellular glucose concentration from 2.8 mM to 25 mM. (c) TIRF images of INS1 cells transiently expressing Ins-SNAP labeled with TMR-Star (Ins-SNAP^TMR^) and Syntaxin1-GFP (Stx1-GFP). (d-e) Mean cumulative percentage of Ins-SNAP^TMR^/Stx1-GFP^+^ objects in living INS-1 cells exposed to 2.8 mM glucose, treated with the indicated siRNAs, and imaged by TIRF microscopy (30 frame/sec) for 7.2 sec. Data in (d) were collected from 31 and 32 movies imaging in total 6.43 x10^2^ and 4.67×10^2^ Ins-SNAP^TMR-Star+^ objects in cells treated with *scrambled* or *vil* siRNAs, respectively. Unpaired two-tailed t-test; p < 0.0001. Data in (e) were collected from 10, 10 and 9 movies imaging in total 4,64×10^2^, 5,14×10^2^ or 5,27×10^2^ Ins-SNAP^TMR^ objects in cells treated with *scrambled siRNA + CMV-Mock, Vil* siRNA + *CMV-Mock or Vil* siRNA + *CMV-Vil,* respectively. Unpaired two-tailed t-test; p = 0.0011. p-values: * ≤0.05; ** ≤0.01; *** ≤0.001; **** ≤0.0001. (f) Percentage of nearly immobile objects (D∼<10^−4^ µm^2^/sec).

### *Villin* depletion enhances the immunodetection of Snap25

Next, we examined the expression and distribution of proteins involved in the docking of SGs to the plasma membrane. Immunoblotting indicated that villin depletion did not alter the levels of the t-SNAREs Stx1, Stx4, Snap25a, Snap25b, or the SNARE interactor Munc-18 (Fig. 2a). Confocal imaging of cells transfected with villin-targeting shRNA within the pCLIP reporter plasmid, albeit less efficient than siRNA-mediated silencing (Fig. 2a–b), allowed the identification of transfected cells and revealed a marked increase in SNAP25 immunoreactivity in villin-depleted INS-1 cells, whereas Stx1 staining was unchanged (Fig. 2c–d). Conversely, overexpression of villin-GFP significantly reduced SNAP25 immunoreactivity (Fig. 2e). A plausible explanation for these observations is that proximity of villin to SNAP25 hinders the immunodetection of the latter. Nevertheless, even upon co-overexpression of villin-GFP and SNAP25-mCherry, their colocalization appeared restricted to the cell cortex (Extended Data Fig. 2 a,b).

**Figure 2.**
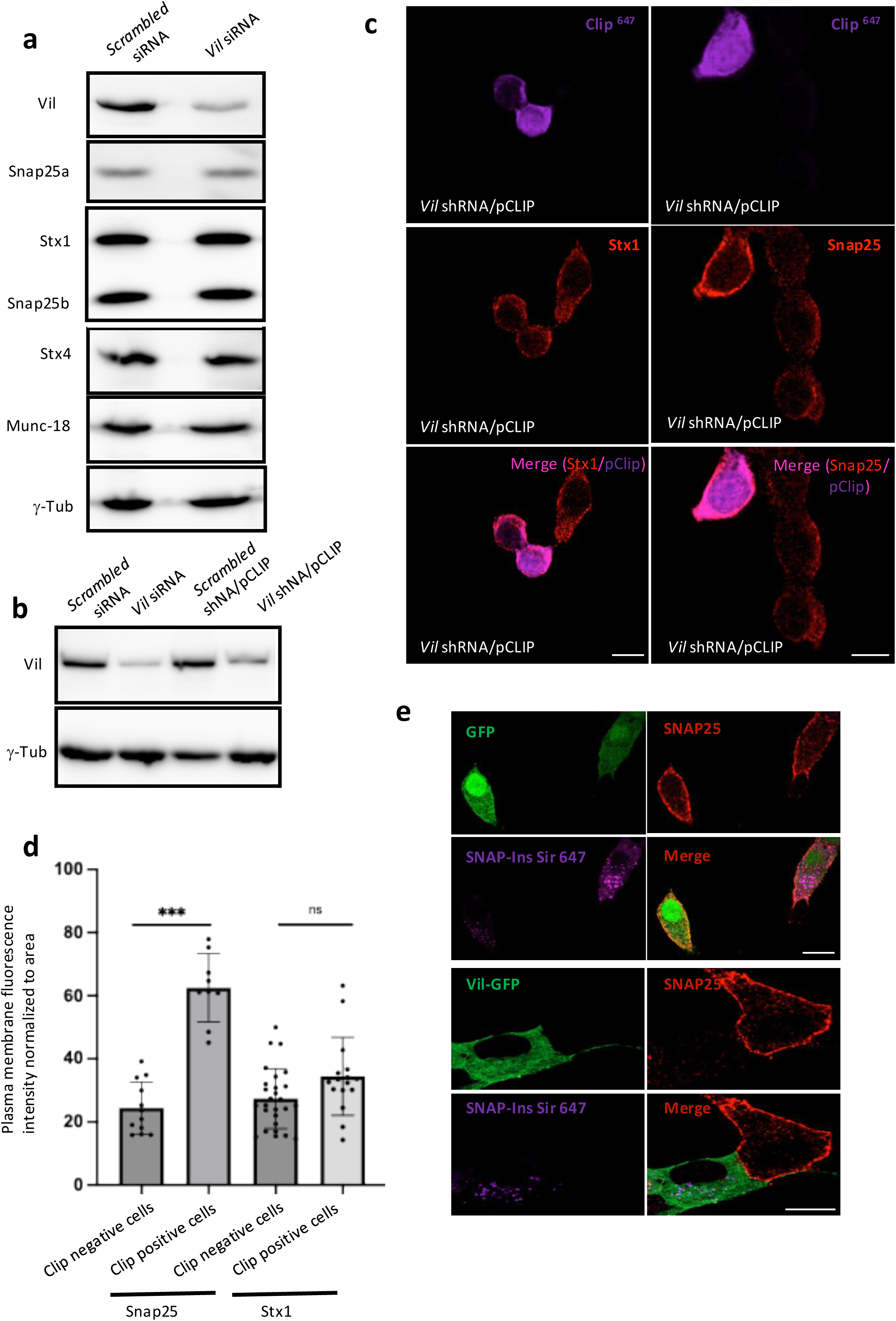
*Vil* depletion enhances SNAP25 immunoreactivity in INS-1 cells. (a-b) Western blotting for villin, SNAP25a, SNAP25b, STX1, STX4, MUNC18 and γ-tubulin in extracts of INS-1 cells treated with *scrambled* or *Vil* siRNAs or with short hairpin in the pCLIP reporter plasmid (b). (c) Confocal microscopy images of control versus *villin*-depleted INS-1 cells immunostained for SNAP25 or STX1. (d) Mean immunofluorescence intensity for SNAP25 and STX1 in INS-1 cells treated with *scrambled* shRNA or *Vil* sihRNA. Each dot corresponds to the measurement in one cell. Unpaired two-tailed t-test; n = 6; mean difference = 38.5 ± 5.0 (SEM); p = 0.0006. Scale bars, 10 µm. p-values: p-values: * ≤0.05; *** ≤0.001. (e) Confocal microscopy images of INS-1 cells expressing Snap-Ins labeled with Sir 647 (SNAP-Ins^Sir 647^) co-transfected either with GFP or Vil-GFP immunostained for SNAP25.

### Villin binds to Snap25

To investigate further the relationship between villin and SNAP25b, we resorted to microscale thermophoresis, which is a sensitive *in vitro* approach to query protein-protein interactions (Magnez, et al, 2017). Extracts of INS-1 cells (Extended Data, Fig. 3a) expressing mCherry or Snap25b-mCherry were incubated with extracts of INS-1 cells overexpressing villin or Stx1 (Fig. 3a). In both cases the temperature-induced fluorescent profile of mCherry did not change, whereas that of SNAP25b-mCherry displayed a comparable shift upon incubation with extracts enriched for its *bonafide* interactor Stx1 or villin. Conversely, the fluorescence profiles of phosphoserine phosphatase (PSPH)– and Myoglobulin-mCherry expressing INS-1 cell extracts were unaffected by exposure to extracts of villin-enriched INS-1 cells (Extended Data, Fig. 3b), pointing to the specific binding of SNAP25b to villin.

**Figure 3.**
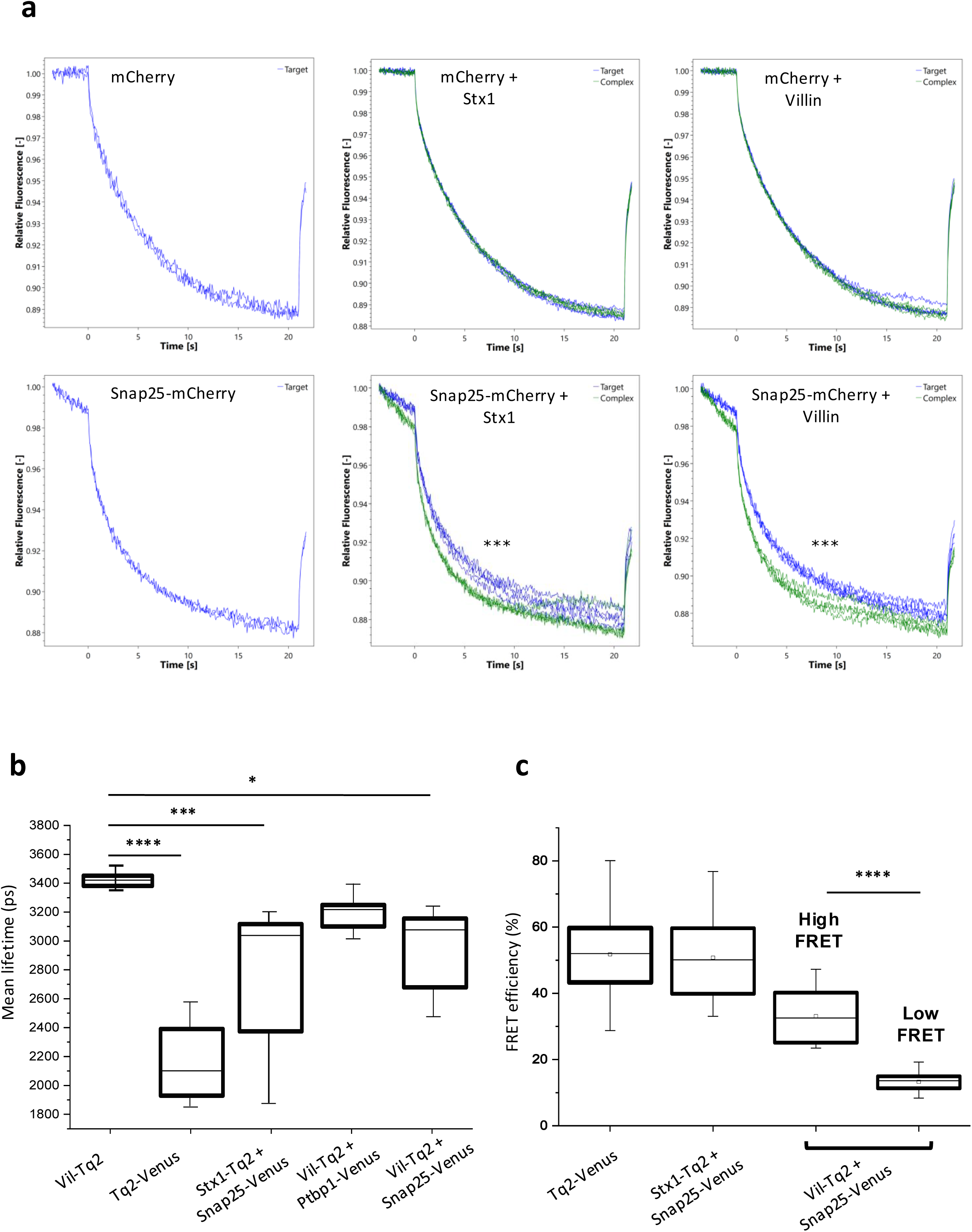
Villin directly interacts with SNAP25. (a) Representative micro scale thermophoresis plots of the fluorescence for mCherry upon mixture of extracts of INS-1 cells transiently transfected with mCherry, Snap25-mCherry with extracts of INS-1 cells transfected with villin or Stx1, as indicated. The capillaries were heated for 20 sec and the fluorescence intensity decay of the reporter proteins (mCherry fluorescence) was analyzed with the NanoTemper software. The green and the blue traces display the fluorescence of the unbound and the bound fractions of the reporter proteins, respectively. Unpaired two-tailed t-test; n = 3; p≤ 0.001. (b) Average lifetime of the FRET signal in INS-1 cells transiently transfected with the indicated constructs as measured by fitting the decay of vil-Tq2 with a single exponential function with a time constant of 3.5 ns (c) FRET efficiency in INS-1 cells expressing the indicated constructs. Results in (a) are from **3** and in (b-c) are from 7-15 independent experiments. p-values: * ≤0.05; *** ≤0.001; **** ≤0.0001.

Further evidence for the interaction of villin with SNAP25b was obtained through FLIM-FRET analyses in INS-1 cells. In this case, villin and Stx1 tagged with monomeric Turquoise2 (Tq2) were co-expressed as donors in various combinations with SNAP25b or Ptbp1 as a negative control fused to fluorescent Venus as acceptors (Extended Data Fig. 3c). In our setting the donor lowest lifetime intensity, indicative of donor-acceptor interaction, was set by expressing a tandem construct in which Tq2 and Venus were separated only by a short linker (Fig. 3b and Extended Data Fig. 3c), whereas the highest lifetime intensity was measured by expressing vil-Tq2 alone (Fig. 3b and Extended Data Fig. 3d). Expression of SNAP25b-Venus with Vil-Tq2 or Stx1-Tq2 quenched the donor’s mean lifetime by 466±153ps (p=0,038) or 641±137,64 ps (p=0,0002), respectively (Fig. 3b and Extended Data Fig. 3d), whereas the quenching of vil-Tq2 mean lifetime by Ptbp1-Venus was not significant (299±129ps; p=0,257) (Fig. 3b). The binomial distribution of the FRET signal between villin-Tq2 and SNAP25-mCherry is indicative of a high and a low affinity interaction (Fig. 3c) and compatible with villin being present in distinct conformations located at different distances from SNAP25 (Extended Data Fig. 3e).

### The N-terminal region of villin binds to the coiled-coil domain 2 of SNAP25 in a glucose dependent manner

The binding of villin to SNAP25 was further tested by co-immunoprecipitations followed by immunoblotting with antibodies directed against different domains of the two proteins (Fig. 4a). To circumvent interpretative difficulties due to the overlapping electrophoretic mobility of SNAP25 and the immunoglobulin light chains, these tests were also performed using extracts of INS-1 cells expressing SNAP25-mCherry. As shown in Fig. 4b, the antibody directed against the SNAP25 D1 coiled-coil domain immunoprecipitated untagged, overexpressed villin, and vice versa, the antibody directed against the villin C-terminal region pulled down SNAP25-mCherry (Extended Data Fig. 4a). To map the region of SNAP25b binding to villin, its D1 (19-81 aa) and D2 (140-202 aa) coiled-coil domains were independently expressed in bacteria as GST-tagged proteins (Fig. 4a) for pull-down assays from extracts of INS-1 cells overexpressing villin. Only SNAP25-D2 bound to the N-terminal portion of villin, which was recovered as a fragment (vil-NTF) (Fig. 4c-d and Extended Data Fig. 4b). Likewise, recombinant GST-villin (1-250) pulled-down endogenous SNAP25 (Fig. 4e), and specifically SNAP25-D2, but not SNAP25-D1 (Extended Data Fig. 4c), as confirmed by mass spectrometry of the recovered peptides excised of their respective GST tags (Extended Data Fig. 4c). Taken together, these data show that the N-terminal region of villin, which contains the two major Ca^2+^ sensitive sites of the protein (Hesterberg and Weber, 1983), directly interacts with the C-terminal D2 domain of SNAP25. Notably, the recovery of villin with SNAP25 from INS-1 cells stimulated with 25 mM glucose was only 25% compared to resting INS-1 cells (Fig. 4f), suggesting a scenario whereby glucose-induced dissociation of the villin-SNAP25 complex enhances access of SGs to t-SNAREs for docking and exocytosis.

**Figure 4.**
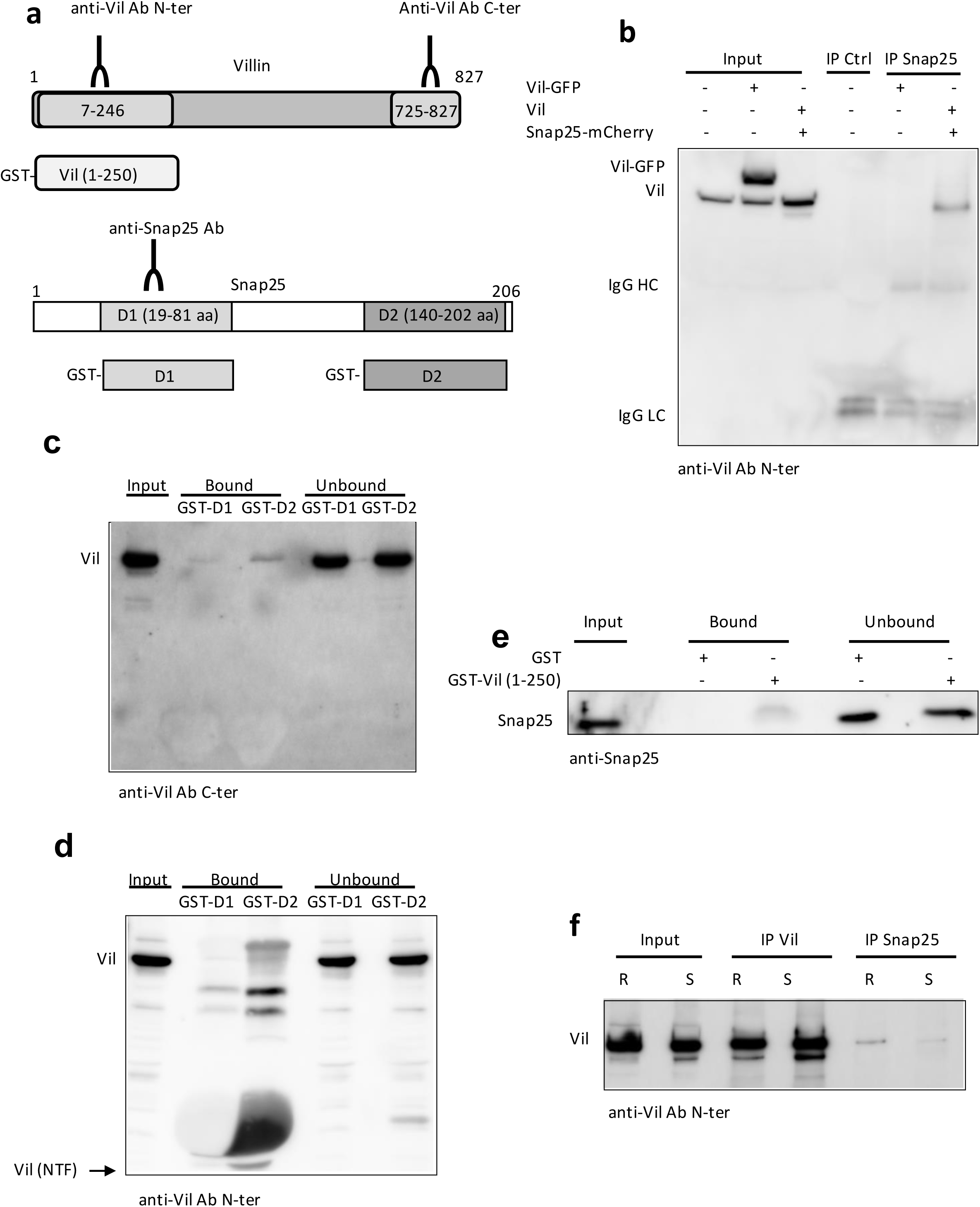
SNAP25 coiled-coil domain D2 interacts with villin N-terminus in a glucose-regulated fashion. (a) Schematic representation of the epitope region recognized by commercial anti-villin(a) and – (b), anti-SNAP25 antibodies and the recombinant GST-villin (1-250), GST-D1 and GST-D2 constructs. (b) Western blots for villin and SNAP25 on immunoprecipitates with control or anti-SNAP25 antibodies from extracts of INS-1 cells transfected with either non-tagged villin, villin-GFP or SNAP25-mCherry. (c-d) Western blot with (c) anti-villin(a) or (d) anti-Viilin(b) antibodies on GST-D1 or GST-D2 pull-down fractions from extracts of INS-1 cells transfected with untagged-villin. (e) Western blot for SNAP25 on GST or GST-Vil (1-250) pull-down fractions from extracts of INS-1 cells. (f) Western blot for villin immunoprecipitated with anti-villin(a) or anti-SNAP25 antibodies from extracts of resting (R, 2.8 mM glucose) or stimulated (S, 25 mM glucose) INS-1 cells co-transfected with non-tagged villin and SNAP25-mCherry.

### Stabilization of ICA512-CCF enhances the levels of PY-Stat5 and villin in INS-1 cells

Our previous studies showed that ICA512 positively regulates villin mRNA and protein levels in insulinoma cells and mouse islets (Mziaut et al, 2016). The close connection between ICA512 and villin was corroborated by evidence that tamoxifen treatment of human EndoC-βH3 β-like cells, which inhibits their proliferation and enhances their insulin content and GSIS (Benazra et al, 2015), correlates with the concomitant upregulation of ICA512 and villin mRNA and protein levels (Extended Data Fig. 5a-d). These data, however, do not explain how ICA512 modulates villin expression. We wondered in particular whether this regulation depends on the ICA512 retrograde signaling elicited with the generation of ICA512-CCF by Ca^2+^/calpain1-induced cleavage of its cytosolic tail upon SG exocytosis (Ort et al, 2001; Trajkovski et al, 2004; Mziaut et al, 2006). Answering this question, however, had been hindered by the inability to detect ICA512-CCF in primary islet β-cells. On the other hand, studies in ICA512 transfected HEK293T cells showed that replacement of lysine 609 with valine (K/V) at the very N-terminus of ICA512-CCF stabilized this calpain-generated short-lived substrate of the Arg/N-end rule pathway, conceivably by preventing its ubiquitination and degradation (Piatkov et al, 2014). Supporting this hypothesis, treatment with proteasome inhibitors epoxomicin, and especially MG-132, increased ICA512-CCF levels in both INS-1 and MIN6 insulinoma cells (Extended Data Fig. 6a-b). Therefore, we compared the western blot profiles of control INS-1 cells with that of INS-1 cells overexpressing human ICA512^wt^ or ICA512^K609V^ tagged with HA. As shown in Fig. 5a-c, the levels of pro-ICA512, ICA512-TMF and ICA512-CCF were greatly enhanced in cells expressing the ICA512^K609V^-HA variant. Moreover, in these cells tyrosine phosphorylated STAT5 (PY-STAT5) and villin were also upregulated (Fig. 5d-e), consistent with stabilization of ICA512-CCF extending STAT5 activity (Mziaut et al, 2006) and driving ICA512’s own expression as well as that of villin. Mass-spectrometry confirmed that the tryptic fragment of the top ICA512-CCF species immunoprecipitated with an anti-HA antibody from ICA512^K609V^-HA INS-1 cells (Fig. 5f) started at leucine 612 (Fig. 5g), adjacent to the most N-terminal calpain-cleavage site located between residues 608 and 609 (Ort et al, 2001).

**Figure 5.**
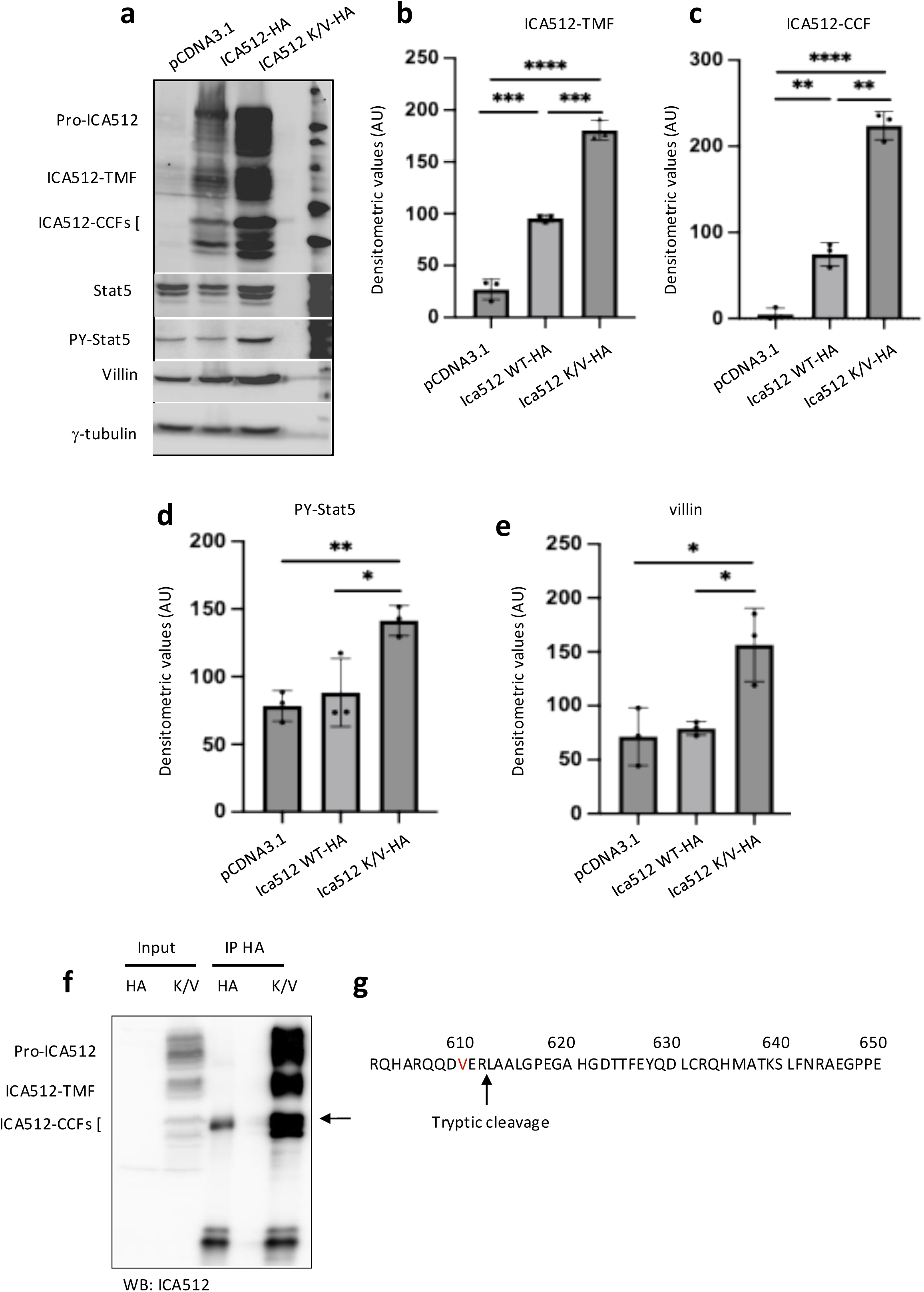
Expression of *ICA512^K609V^* increases the levels of PY-STAT5 and villin. (a) Western blot for ICA512, STAT5, PY-STAT5, villin and γ-tubulin on extracts of INS-1 cells transfected with HA, ICA512-HA or ICA512 K/V and treated with 20 nM growth-hormone in 11 mM glucose for 18 hours. (b-e) Quantification of ICA512-TMF, ICA512-CCF, PY-STAT5 and villin are from at least 3 independent experiments as in (a). (f) Western blot for ICA512 on immunoprecipitates with anti-HA antibodies from extracts of INS-1 cells transfected with HA or human ICA512-K609V (K/V). (g) Primary amino acid sequence of human ICA512 cytosolic fragment (residues 601-950) highlighting the fragment obtained by tryptic cleavage at leucine 612 adjacent to lysine 609 also **shown in table 1**. Statistics in b-e are from unpaired two-tailed t-test: p-values: * ≤0.05; ** ≤0.01. *** ≤0.001.

**Table 1.**
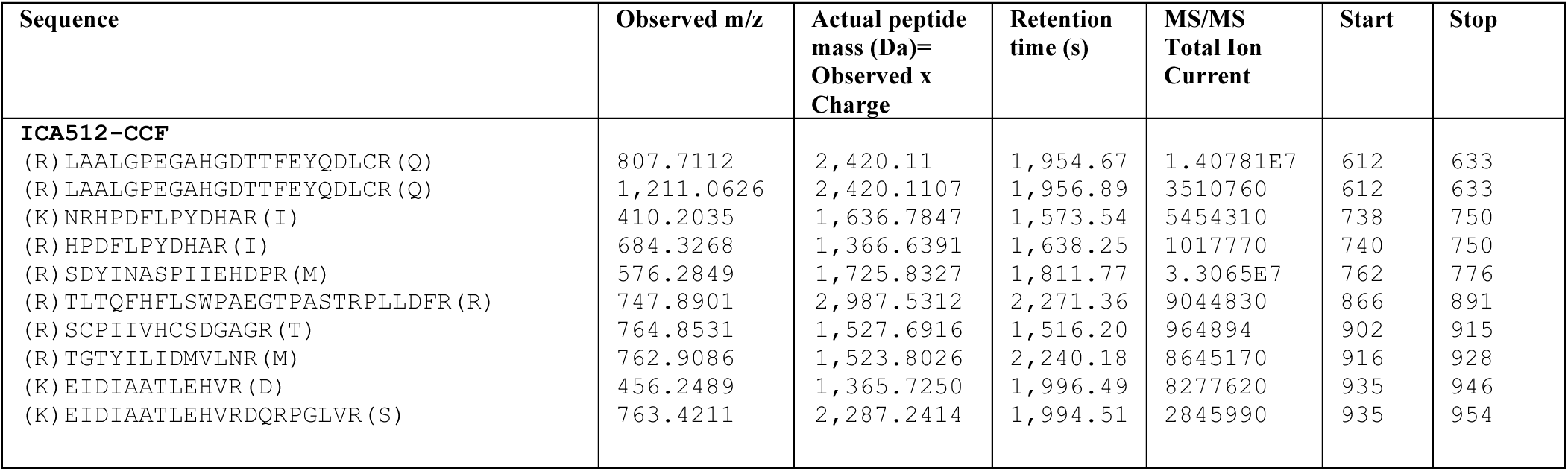
Table showing the amino-acid sequence, observed monoisotopic m/z, average peptide mass, retention time and the location in the protein.

### *ICA512^K609V^* mice display lower body weight and improved insulin sensitivity

To assess whether ICA512-CCF signaling is physiologically relevant, we then generated knock-in *ICA512^K609V^* mice and first analyzed their metabolic profile. Homozygous female, but not males, *ICA512^K609V^* mice displayed a reduced body weight compared to control littermates (females: 18,8±1,2g vs. 21,6±1,4 g; males: 26,1± 0.6 g vs. 27,7±1.8 g) (Fig. 6a-b). Intraperitoneal glucose tolerance tests did not reveal changes in the glycemic profile of *ICA512^K609V^* mice (Fig. 6c and Extended Data 7a), but females, unlike males (Extended Data 7b), showed improved insulin sensitivity compared to controls (Fig. 6d).

**Figure 6.**
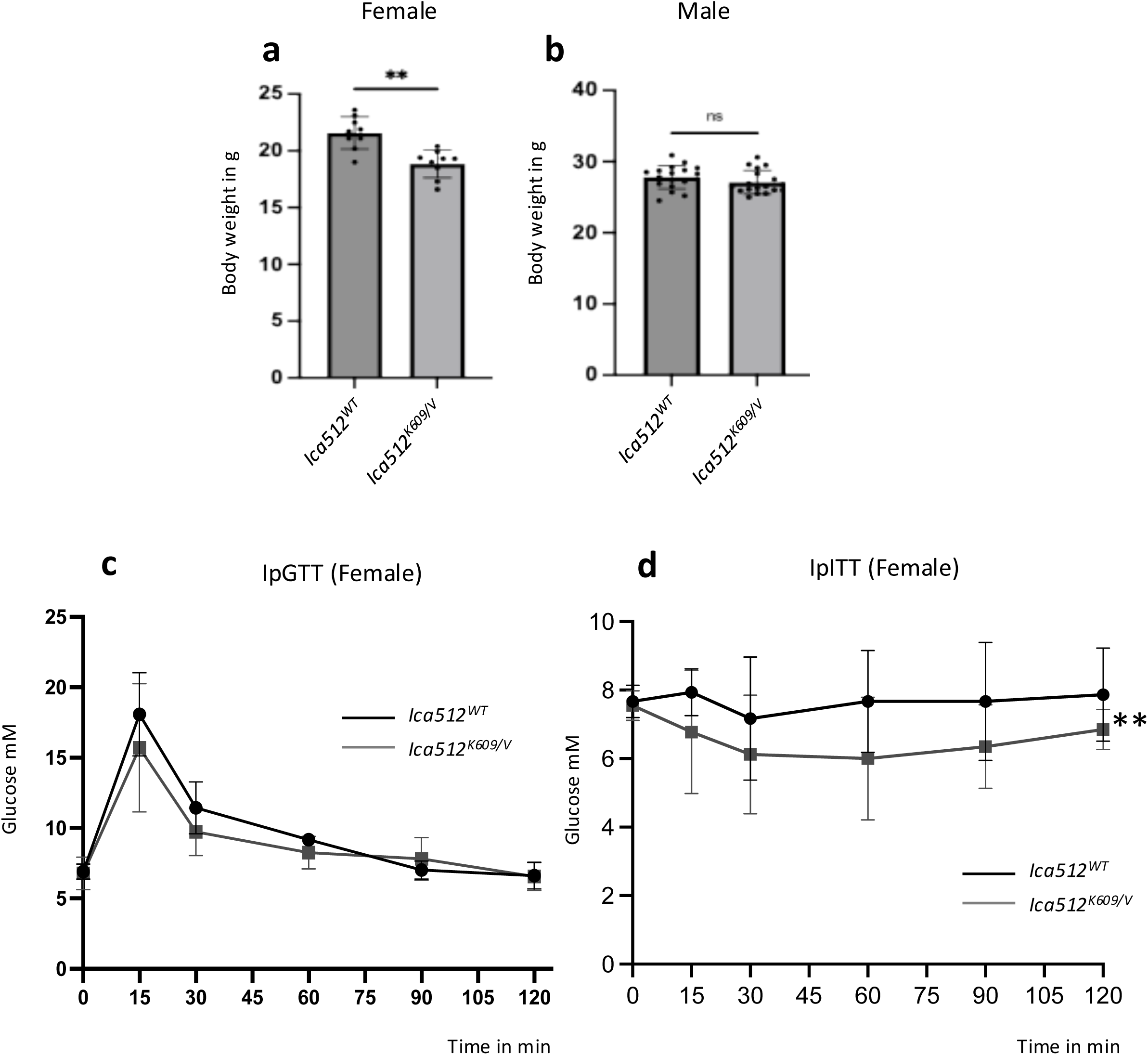
Reduced body weight, improvement in IpITT without alterations in IpGTT in female *Ica512^K609/V^* mice. (a-b) Body weight of 13-15 week old female (a) and 13-14 week old (b) male *Ica512^wt^* and *Ica512^K609/V^* mice. Two-tailed t-test; n = 9 pairs; t(8) = 3.75; p = 0.0056. (c) IpGTT on 6 *Ica512^wt^* and 4 *Ica512^K609/V^* female mice. (d) ipITT on 3 *Ica512^wt^* and 5 *Ica512^K609/V^* female mice. For ipITT, we use coupled test (series of data time dependant) for difference of responses relative to the point t=0 as described in material and methods

### Villin expression and insulin SG stores are increased in *ICA512^K609V^* mouse islets

Moving the analysis to *ICA512^K609V^* mouse islets, their *ICA512* mRNA expression was elevated when cultured in 3.3 mM glucose (Resting, R), but unlike in islets of control littermates, it did not increase further upon costimulation with 16.7 mM glucose (high glucose, HG) and 20 nM growth hormone (GH) for 18 hours (Fig. 7a). *Villin* mRNA levels in resting islets of *ICA512^K609V^* and control mice were comparable, but increased instead in co-stimulated islets of the mutant mice (Fig. 7b). Western blotting of *ICA512^K609V^* islet extracts enabled for the first time to detect the endogenous occurrence of ICA512-CCF in primary islet cells. As expected, its levels as well as those of its precursors pro-ICA512 and ICA512-TMF, two hallmarks of increased SG biogenesis, were enhanced by HG (Fig. 7c and d). In *ICA512^K609V^* islets we could also verify that stabilization of ICA512-CCF by a single amino acid replacement correlated with elevated levels of PY-STAT5 upon stimulation with HG and GH, but not with GH alone, presumably due to the short half-life of PY-STAT5 (Yu et al, 2000). Accordingly, the temporally restricted synergy between glucose and GH signals through the interaction of ICA512-CCF with Y-STAT55 was manifested by the lowering of their expression levels upon removal of both stimuli for 14 hours after co-stimulation for 4 hours (Fig. 7c, lane 10). Similar, albeit not identical, conclusions could be drawn about PY-STAT3 levels being increased in *ICA512^K609V^* mouse islets (Extended Data 8a). Moreover, stabilization of ICA512 retrograde signaling resulted in the increased expression of villin upon stimulation of islets with glucose, either alone or together with GH (Fig. 7c and e), as well as in the upregulation of the insulin SG stores, as shown by morphometry of electron microscopy images (Fig. 7f-g) and measurement of the insulin content (Fig. 7h).

**Figure 7.**
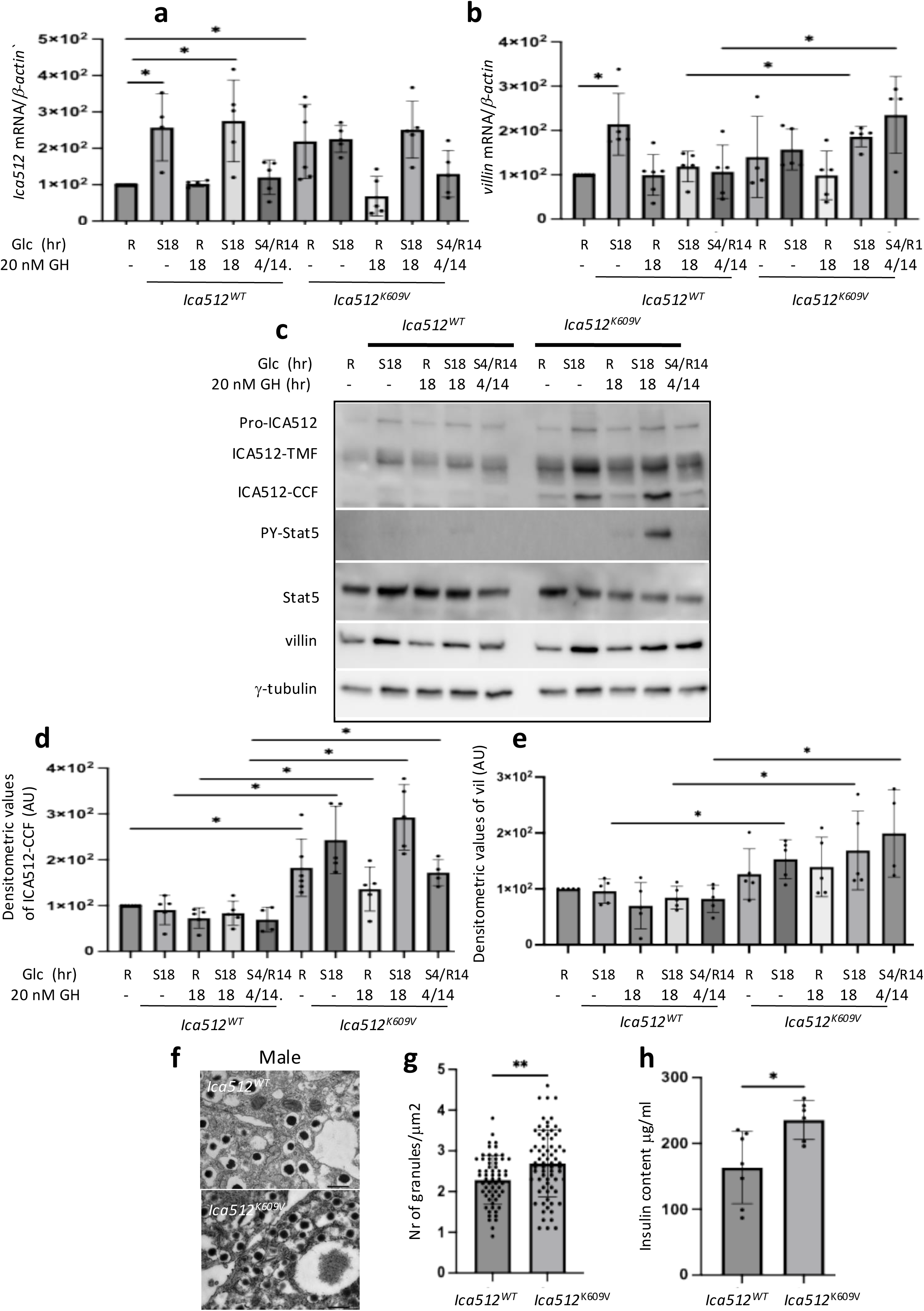
Enhanced STAT5 phosphorylation in correlation with increased insulin content and calcium signaling in *Ica512^K609/V^* mice. (a-b) q-PCR analysis of ICA512 (a) and villin mRNA (b) from mouse pancreatic islets incubated 18 hrs either in resting conditions or in presence of 16,7 mM glucose and 20 nM growth hormone. (c) Western Blot analysis of islets isolated from or *Ica512^w^*^t^ and *Ica512^K609/V^* mice treated as indicated and the blots probed for ICA512, PY-STAT5, STAT5, γ-tubulin and villin antibodies. (d-e) Quantification by densitometric analysis of the proteins as detected in (c). Experiments in (a-e) are at least from 4 independent experiments. Statistics performed using paired two-tailed t-test: p-values: * ≤0.05; ** ≤0.01. (f-g) electron microscopy images (f) and quantification of granule density (g) and insulin content (h) in islets isolated from *Ica512^wt^* and *Ica512^K609/V^* mice. Unpaired two-tailed t-tests. For (g): t(130) = 3.17, p = 0.0019. For (h): t(11) = 2.86, p = 0.0155.

### *ICA512^K609V^* mice islets display improved glucose responsiveness

islets, in which stabilization of ICA512-CCF through a single-point mutation leads to the overexpression of both endogenous ICA512 and villin as well as increases the insulin stores. As shown in Fig. 8a, the first and second phase of GSIS of islets from *ICA512^K609V^* and control mice were comparable. Likewise, no differences in insulin release were measured upon maximal stimulation with 55 mM KCl (high potassium, HK). Yet, the basal insulin release of *ICA512^K609V^* islets was reduced (Fig. 8b), while their stimulation index (stimulated insulin secretion/basal insulin release) in response to 16.7 mM glucose alone (Fig. 8c) or together with HK (Fig. 8d) was enhanced. Further changes were revealed by Ca^2+^ imaging. In glucose stimulated *ICA512^K609V^* islets, both the amplitude (Fig. 8e) and area under the curve (Fig. 8f) of the Ca^2+^ fluorescence intensity were increased, while its onset and rise times were faster (Fig. 8g-h). The rising time of the Ca^2+^ signal was also shorter upon stimulation with HK, while the onset time was unchanged (Fig. 8i-j). The GSIS of male *ICA512^K609V^* islets (Extended Data Fig. 9a), as well as the amplitude, rise time and area under the curve of the glucose induced Ca^2+^ signals were unchanged (Extended data Fig. 9d-e and 9g). However, the onset of their Ca^2+^ fluorescence was faster (Extended data Fig. 9f). Upon stimulation with HK, the onset time of the Ca^2+^ signal in male *ICA512^K609V^* islets was unchanged, but the rise time was reduced (Extended data Fig. 8h-i). Taken together, these results indicate that stabilization of ICA512 retrograde signaling enhances β-cell responsiveness to glucose, likely through downstream regulation of villin expression and the interaction of the latter with SNAP25. Therefore, we finally asked whether the improved glucose responsiveness of ICA512^K609/V^ β-cells might result from altered interaction between villin and SNAP25. To address this question, we performed proximity ligation assays (PLA) on primary islets isolated from *Ica512*^wt^ and Ica512^K609V^ mice. As shown in Fig. 8 k-l, the number of villin–SNAP25 PLA puncta per DAPI⁺ cell was significantly higher in ICA512^K609V^ islets compared with wild type (8.2 vs 2.5 ±, 1,43, p=0.0001), indicating an increased association between the two proteins. The enhanced villin–SNAP25 interaction likely accounts, at least in part, for the improved exocytotic behavior of ICA512^K609V^ β-cells characterized by reduced basal insulin secretion but accelerated glucose-induced Ca²⁺ signaling and insulin release.

**Figure 8.**
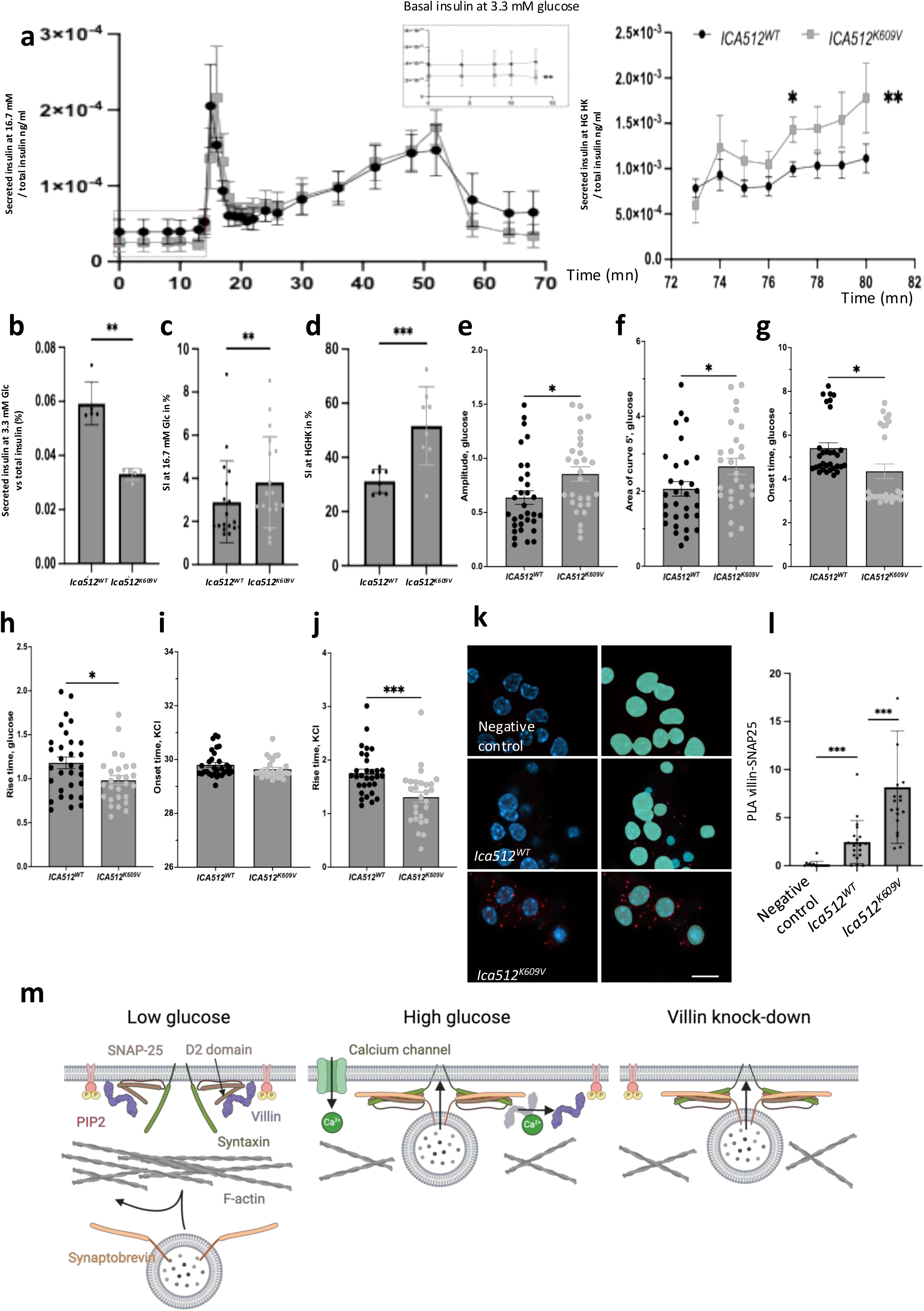
(a) Dynamic GSIS on islets isolated from *Ica512^wt^* and *Ica512^K609/V^* mice. For clarity, the data are presented in three separate graphs showing: secreted insulin at basal and first-phase glucose stimulation (left panel), second-phase GSIS (middle panel), and insulin secretion in response to high glucose plus KCl (HGHK, right panel). plots (b-d) representing secreted insulin vs total at 3.3 mM glucose (b), stimulatory index (SI) at 16.7 mM glucose (c) or SI in response to 16.7 mM glucose combined with 55 mM KCl (d). Calcium imaging of glucose– and HKCl-stimulated islets from *Ica512^wt^* and ICA512^K609/V^ mice. (e–f) Increased amplitude and area under the curve (AUC) of glucose-induced Ca²⁺ transients in *Ica512^wt^* and ICA512^K609/V^ mice islets. (g–h) Reduced onset and rise times of Ca²⁺ responses to glucose stimulation. (i–j) Shortened rise time, but unchanged onset, of Ca²⁺ responses upon HKCl stimulation. Statistical significance was determined using a two-tailed t-test; p-values: * ≤0.05; ** ≤0.01;. Proximity ligation assay (PLA) analysis of villin–SNAP25 interaction in primary islets. (k) Representative PLA images showing raw villin–SNAP25 puncta (red) per DAPI⁺ nucleus (blue) in *Ica512^wt^* and ICA512^K609/V^ islets compared with wild type (left panels). Nuclei and villin–SNAP25 puncta were automatically detected and quantified using Arivis Pro /Zeiss (right panels). Scale bars, 10 µm. (l) Quantification of villin–SNAP25 PLA puncta per DAPI⁺ cell. Data are shown as mean ± s.e.m. from n = 2 independent experiments, with ∼10 images analyzed per experiment. Statistical significance was determined using a two-tailed t-test; p-values: * ≤0.05; ** ≤0.01. (m) Villin is recruited to the plasma membrane through PIP₂, positioning it near SNAP25 within the SNARE complex. At low glucose, villin interacts with SNAP25, likely masking or sterically hindering its docking-competent conformation, thereby limiting secretory granule (SG) docking. Depletion of villin increases SG docking even in the absence of stimulation. Upon glucose stimulation, the resulting rise in intracellular Ca²⁺ correlates with villin’s dissociation from SNAP25 and activation of its severing function, enabling SGs to access their docking sites.

## Discussion

In this study, we further explored the role of the actin modifier villin in pancreatic β-cells. This gelsolin paralog bundles cortical F-actin at low Ca²⁺ concentrations and severs or caps it when Ca²⁺ levels rise (Khurana and George, 2008), as during glucose stimulation. We previously reported that villin expression is positively regulated by the SG cargo ICA512/PTPRN and that its depletion in INS-1 cells increases basal insulin release, likely due to reduced cortical actin bundling and thus enhanced SG mobility (Mziaut et al, 2016). However, modeling predicted that increased SG mobility alone could not account for enhanced basal insulin release unless the number of accessible docking sites for exocytosis is also increased. Supporting this hypothesis, we identified the N-terminal domain of villin as a binding partner of the D2 domain of SNAP25, a t-SNARE essential for insulin SG exocytosis. This interaction was reduced upon glucose stimulation, suggesting that its dynamics regulates the pool of SNAP25 molecules accessible for SG docking. Comparable actin–SNARE regulatory mechanisms have been described for gelsolin–Syntaxin4 in MIN6 cells (Kalwat et al, 2014). Consistently, both villin and gelsolin are upregulated by prolonged glucose exposure (Waanders et al, 2009), and low gelsolin expression correlates with impaired GSIS (Tomas et al, 2006) in MIN6 clones refractory to GSIS, in line with our observation that villin depletion compromises stimulus-secretion coupling (Mziaut et al, 2016). Based on the data presented here, we propose a model whereby villin–SNAP25 interaction at low glucose helps to maintain a fraction of docking sites in a refractory state, while villin–SNAP25 dissociation in response to glucose stimulation and Ca²⁺ influx converts these sites into fusion-competent ones. We also found indications that villin undergoes proteolytic modification upon β-cell stimulation, generating an N-terminal fragment which binds to SNAP25. Notably, the generation of a proteolytic villin N-terminal fragment in intestinal cells exposed to different insults has been proposed to promote apoptosis (Wang et al, 2016). It is also interesting to relate these observations to earlier morphological analyses in diabetic models. In Zucker Diabetic Fatty rats, Unger and colleagues reported a marked decrease in the volume density of GLUT2-positive β-cells, accompanied by a reduction in microvillar coverage of the plasma membrane (48% in nondiabetic males versus 37% and 30% in mild and severe diabetes, respectively). Given that β-cell microvilli contain the highest abundance of GLUT2 (Orci et al, 1990), these findings raise the possibility that villin-dependent regulation of microvillar architecture may contribute to impaired glucose sensing under diabetic conditions.

The importance of cytoskeletal regulation for β-cell differentiation and function is underscored by recent developmental studies. Differentiation of human pluripotent stem cells into mature β-like cells remains inefficient, partly due to inadequate cytoskeletal remodeling (d’Amour et al, 2006; Nair et al, 2019). Actin polymerization has been shown to influence endodermal specification (Hogrebe et al, 2020) and the generation of functional islets (Karanth et al, 2021). In tamoxifen-treated EndoC-βH3 cells, which undergo cell-cycle arrest and acquire β-cell characteristics (Benazra et al, 2015), we observed concomitant upregulation of ICA512 and villin, suggesting a coordinated program that links β-cell maturation and SG biogenesis with cytoskeletal adaptation.

To dissect this relationship in vivo, we generated *Ica512^K609/V^* knock-in mice in which a single-point mutation stabilizes the ICA512 cytoplasmic fragment and enhances its retrograde signaling. *Ica512^K609/V^* islets exhibited increased ICA512 and villin expression, expanded insulin SG stores, and heightened responsiveness to glucose. These changes were accompanied by an increased number of villin–SNAP25 complexes, as revealed by proximity ligation assays, indicating that ICA512 stabilization reinforces in parallel the coupling between actin remodeling and SNARE-mediated exocytosis. Accordingly, *Ica512^K609/V^* islets displayed reduced basal secretion but faster Ca²⁺ dynamics and potentiated glucose-evoked insulin release—hallmarks of improved stimulus–secretion coupling. Notably, female *Ica512^K609/V^* mice also displayed lower body weight and improved insulin sensitivity, consistent with the reduction in basal insulin output. Since basal insulin secretion and insulin sensitivity are inversely correlated, this coordinated phenotype suggests that ICA512 stabilization supports a systemic adaptation in which β-cells operate at a lower basal secretory tone while maintaining rapid responsiveness to metabolic demand.

In summary, our findings identify ICA512 as a physiological molecular integrator linking β-cell secretory competence with peripheral insulin sensitivity through regulation of villin and its interaction with SNAP25. We propose that ICA512-mediated control of villin expression dynamically coordinates actin remodeling for the recruitment and fusion of insulin SGs, providing a mechanism by which β-cells fine-tune the ability to secrete insulin in relationship to metabolic needs.

## Material and Methods

### Animals

All animal experiments were conducted in accordance with European Union animal welfare legislation and were approved by the competent authorities (permit numbers TVvG 10/2023 and TVV 40/2020). Experiments were performed in C57BL/6NCrl mice. Animals were housed under a 12 h light/12 h dark cycle at 20–24 °C with 55 ± 10% relative humidity, in individually ventilated cages (IVC; ∼540 cm²), and had ad libitum access to water and a sterilized standard diet (Teklad Global Diet 2018S, Envigo). Unless otherwise indicated, mice used in this study were 10–12 weeks of age; animals used for body weight measurements were 13–15 weeks old. The sex and number of animals used for each experiment are indicated in the figure legends. Animals were allocated to experimental groups based on genotype; no randomization or blinding was performed.

### Culture of mouse islets and insulinoma INS-1 and EndoC-**β**H1cells

Rat INS-1 insulinoma cells were kind gifts from Dr. C. Wollheim (University of Geneva, Switzerland) and grown as described in (Ort et al, 2001). EndoC-βH1 and EndoC-βH3 cells were a gift from Dr. R. Scharfmann (University of Paris Descartes, France) and grown as described (Benazra et al, 2015).

### cDNA constructs and siRNA oligonucleotides

The following plasmids pEGFP-N1, Turquoise, mCherry and Venus were from Clonetech (Foster City, CA). The plasmids used to induce the expression of human *Ica512-GFP* and *insulin-SNAP* have been described elsewhere (Trajkovski et al, 2004), (Ivanova et al, 2013). The mouse *villin* cDNAs full length were cloned as an *EcoRI*-*AgeI* or *EcoRI-Not1* inserts into pEGFP-N1 and pGEX6P1 respectively, whereas vil-NTF (aa:1-250) cDNA was cloned as *EcoRI*-*AgeI* insert into pGEX6P1 using the oligonucleotides described in Mziaut et al, 2016. The cDNAs of mouse or rat *SNAP25*, mouse *syntaxin1,* human *myog,* rat *PSPH and rat PTBP1* were inserted in the reporter plasmids using oligonucleotides described in Extended Data Table 2 and then sub-cloned in various reporter plasmids using the restrictions HindIII-AgeI.

For the silencing of villin, we used synthetic small interfering RNA (siRNA) oligonucleotides targeting rat *villin* as previously described (Mizaut et al, 2016). Cells were harvested 4 days after transfection and gene knockdown was verified by real-time PCR and Western blot as described in (Knoch et al, 2004).

### Islet isolation and dispersion

Pancreatic islets were isolated by collagenase perfusion of the pancreas as described (Li et al., 2009). Briefly, the pancreas was inflated via bile duct injection with collagenase P, digested at 37°C, and islets were purified by density-gradient separation and hand-picked. Islets were recovered overnight and then dispersed into single cells by incubation with Accutase (or 0.05% trypsin-EDTA) at 37 °C for 5–10 min with gentle trituration,

### Cell extracts and western blotting

INS-1 cells were washed with ice-cold PBS and extracted in lysis buffer (10 mM Tris–HCl, pH 8.0, 140 mM NaCl, 1% Triton X-100, 1 mM EDTA, 1 mM phenylmethylsulphonyl fluoride, 1% phosphatase inhibitors [Calbiochem/Merck Millipore], and 1% protease inhibitor mixture [Sigma, St. Louis, MO]) at 4 °C. Aliquots of 20 µg were separated by SDS-PAGE, and western blotting was performed as described (Mziaut et al, 2008). The source, species and dilutions of antibodies used for immunoblotting are listed in Extended Data Table 1.

### Immunoprecipitation

INS-1 cells were transfected as indicated in Fig. legends were harvested 4 days post-transfection. Briefly, cells were lysed in RIPA buffer (50mM Tris-HCl pH8, 150mM NaCl, 0.1% SDS, 1% NP-40, 0.5% sodium deoxycholate) with Protease and Phosphatase inhibitors. Cells were centrifuged for 10 min at 10 000 rpm and the supernatant was precleared with protein G-Sepharose for 1 hr at 4°C and then incubated with the corresponding antibodies overnight at 4°C with horizontal rotation. The complex antibody-antigen was pulled down with 30 ul of 50% slurry protein G-Sepahrose, washed 4 times with RIPA buffer and 1 time with PBS before being boiled and analyzed by western blotting.

### GST pull-down assays

The coiled-coil domains D1 (aa 19–81) and D2 (aa 140–202) of rat Snap25b, as well as the N-terminal region of villin (aa 1–250), were cloned into pGEX-6P-1 vectors and expressed in *E. coli* as GST fusion proteins. After purification on glutathione–Sepharose beads, equivalent amounts of GST or GST-fusion proteins were incubated with lysates of INS-1 cells for 2 h at 4 °C. Beads were washed extensively, and bound proteins were eluted, separated by SDS–PAGE, and analyzed by immunoblotting using anti-villin and anti-Snap25 antibodies. For mass spectrometry, bound proteins were eluted with reduced glutathione and subjected to in-gel trypsin digestion followed by LC–MS/MS analysis. Quantification of band intensity was performed by densitometry and normalized to input levels.

### Confocal and super resolution microscopy

INS-1 and EndoC-βH1 cells transfected as indicated in Fig. legends were immunolabeled as previously described (Trajkovski et al, 2004). In some instances, cells were incubated in resting buffer with 2.8 mM glucose or stimulated with 25 mM glucose and 55 mM KCl for 1.5 h before being fixed. EndoC-βH1 cells were cultured for 3 weeks before induction with TAM as described in (Benazra et al, 2015). For immunofluorescent staining, cells were fixed in 4% paraformaldehyde, blocked with 5% goat serum in PBS for 1 h at room temperature. The cells were incubated overnight at 4°C with the primary antibodies diluted as indicated in Extended Data Table 1. The next day, samples were washed and incubated with the secondary antibodies for 2 h at room temperature. Nuclei were counterstained with 4′,6-diamidino-2-phenylindole (Sigma), and coverslips were mounted with Mowiol (Calbiochem/Merck Millipore). Images were acquired at room temperature with an inverted confocal microscope (Zeiss Axiovert 200M) equipped with a Plan-Apochromat × 63 oil objective, numerical aperture 1.4, a Zeiss LSM780 scan head with photomultiplier tubes, and Zeiss LSM 510 AIM software version 4 (Zeiss, Göttingen, Germany). Immunofluorescence intensity was measured using the region of interest in Fiji. High resolution microscopy images were acquired with N-Storm super resolution built on Nikon Ti2 wild field inverted microscope. Images were imported and analyzed with MotionTracking/Kalaimoscope software (Transinsight GmbH, Dresden, Germany), as previously described (Hoboth et al, 2015). For the quantification of SNAP25 puncta staining in control versus villin depleted cells, INS-1 cells were co-transfected with phalloidin to visualize cortical actin. Images were acquired using high resolution spinning disk microscopy and distribution of SNAP25 puncta within 0.6 um of cell contour visualized by phalloidin was measured using MotionTracking/Kalaimoscope software.

### FLIM/FRET

INS-1 cells transfected were cultured in 35 mm petri-dishes (MatTek, MA, USA) for 4 days. FLIM was recorded on live cells using a Zeiss microscope equipped with a 40×1.4 NA water immersion objective. Confocal images of the cells were recorded prior to each FLIM measurement. Fluorescence of Tq2 was excited at 430 nm pulsed with a laser at 80 MHz and detected 570 nM Pixel size was 0.10 um and 512 x 512 pixels and images collected at rate of 1 frame per 6 s. An average of 2000 photons per cell were collected with a photomultiplier tube and processed by a PicoHarp 300 Time-Correlated Single Photon Counting (TCSPC) system (PicoQuant, Berlin, Germany). The fluorescence decay time course fitted a double-exponential function, with time constants of 3.5 ns for Tq2.

### Total internal reflection fluorescence microscopy (TIRFM)

INS-1 cells transfected with insulin-SNAP and stx1-GFP were cultured on coverslips in 6 well plate. On day 4, insulin-SNAP^+^ SGs were labeled as previously described (Ivanova et al, 2013) with the SNAP substrate TMR-Star (New England Biolabs, Ipswich, MA). Prior to imaging, cells grown in an open chamber were incubated in resting media and transferred onto a thermostat-controlled (37 °C) stage. The SGs were visualized as previously described (Mziaut et al, 2016). Automated image analysis was performed with MotionTracking/Kalaimoscope software as previously described (Hoboth et al, 2015). The density of cortical SGs was calculated using ≥38 movies of cells expressing insulin-SNAP and stx1-GFP. Images from different frames were processed post-acquisition using a script derived from the algorithm, called MUltiple SIgnal Classification ALgorithm (MUSICAL) (Agarwal et al, 2016) implemented in motion tracking to achieve high-resolution imaging. The docked granules were defined as granules that remain confined at the plasma membrane for 60 seconds (Omar-Hmead**i** et al, 2018).

### Transmission electron microscopy

Electron microscopy was performed as described in Fava et al. (2012). Briefly, pancreatic islets were fixed in 2.5% glutaraldehyde in 0.1 M cacodylate buffer (pH 7.4), post-fixed in 1% osmium tetroxide, dehydrated in graded ethanol, and embedded in Epon resin. Ultrathin sections (70 nm) were contrasted with uranyl acetate and lead citrate and examined using a Tecnai 12 Biotwin transmission electron microscope (FEI Company, Hillsboro, OR, USA) equipped with a bottom-mounted 2 × 2K F214 CCD camera (TVIPS, Gauting, Germany). Micrographs were taken at 6,800× magnification, and composite images of complete β-cells were acquired using EM Menu 3 software (TVIPS)

### *In silico* modeling of secretory granule dynamics

We developed a spatial β-cell model in which SGs movement is mediated and constrained within the cortical cytoskeleton that is based on experimental data published by Mziaut et al., 2016. The description of the model was detailed in Dehghany et al, 2015. Briefly, in the original model, the number of DSs was assumed to be directly regulated by the glucose concentration. Here, we revised the model so that DS numbers were coupled to the subset of the granule population undergoing exocytosis: Each exocytotic event adds new DSs to the cell surface, increasing the probability of subsequent docking and fusion events. The docking sites were assumed to be removed with a constant rate to ensure homeostasis.

The model adapted the DS number based on the average exocytosis events per minute over the last 20 minutes of the simulation. Specifically, the DS number was determined by the equation 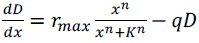: a DS creation rate that depends on recent exocytosis events (normalized to the DS number over the last 20 minutes) minus a DS removal rate.

Abbreviations:

- *D*: docking site number
- *x*: fraction of exocytosed secretory granules (ratio of exocytosis events to total SG number), averaged over the last 20 minutes
- *r_max_*: maximum DS delivery rate
- *K*: half-maximal activation constant
- *n*: assumed steepness of sigmoid function
- *q*: DS removal rate

### Thermophoresis

INS-1 cells seeded in 6 well plates were transfected with 3 ug of plasmids encoding for m-Cherry, SNAP25-mCherry, –Myog-mCherry, PSH-mCherry, villin and Stx1 using Nucleofactor® kit program X. On day 4, cells were washed with PBS, scrapped and centrifuged at 12,000 rpm at 4°C for 5 mn. The cell pellet was resuspended in 150 ul RIPA buffer containing 1x protease inhibitor cocktail. The cell lysate was collected by centrifugation at 10,000 rpm for 10 mn. An aliquot was stored to check for the transfection efficiency by western blotting and the rest was processed for MST measurements. The MST binding assay was performed with Monolith NT.115 instrument (NanoTemper Technologies, Munich, Germany). The cell lysate from each transfection was diluted 5 times in RIPA buffer and a mixture of 20 ul of cell extract containing the reporter mCherry either alone or as fusion protein with SNAP25, myoglobin or PSH was mixed with 20 ul of cell extracts expressing villin or stx1 and incubated at RT for 10 min. The mixture was transferred to capillaries and the fluorescence analyzed using 80% of the MST power.

### Dynamic perfusion

Insulin secretion was performed on islets isolated from male and non-pregnant mice at age 12week old as well as pregnant mice at G13. Briefly, 3 mice/group were sacrificed and 40 islets/mouse were hand-picked, placed into a closed perfusion chamber and analyzed as previously described (Cohrs et al, 2020). After the perfusion experiments, islets were lysed in acid ethanol (2% HCl [37%, 12M] in abs. ethanol) for total insulin content measurements. Samples were kept at −20°C until measured with an Insulin Ultra Sensitive Assay Kit (HTRF from Revvity cat. no. 62IN2PEH).

### In Vitro Imaging of pancreatic islets Cytosolic Free Calcium Dynamics

For in vitro imaging of cytosolic calcium in mouse islets, isolated islets from wt or mutant mice were handpicked after overnight culture and embedded in fibrin gels on coverslips according to a previously published protocol (1). Briefly, fibrin gels were prepared by mixing 5 mL Hanks’ balanced salt solution (HBSS) with 2 mL human thrombin (50 units/mL in HBSS; Sigma-Aldrich), after which ten to fifteen islets were suspended in 2 mL human fibrinogen (10 mg/mL in HBSS; Sigma-Aldrich) and placed individually in the gel to induce fibrinogen polymerization. Subsequently, the gel-embedded islets were loaded with Cal-520® AM (3.33 mM f.c.; AAT Bioquest) working solution in KRBH buffer (137 mM NaCl, 5.36 mM KCl, 0.34 mM Na_2_HPO_4_, 0.81 mM MgSO_4_, 4.17 mM NaHCO_3_, 1.26 mM CaCl_2_, 0.44 mM KH_2_PO_4_, 10 mM HEPES, 0.1% BSA, 3 mM glucose, pH 7.3) and incubated in a cell culture incubator at 37 °C for 30 minutes. Calcium imaging was performed in a custom-made perifusion system using an upright laser scanning microscope (LSM780 NLO; Carl Zeiss, Jena, Germany) as described previously (2; 3). Briefly, coverslips with gel-embedded islets were moved into a temperature-controlled perifusion chamber where the temperature was set at 37°C and the flow rate was set at 100 ml/min. Cal-520® AM was excited at 488 nm and detected at 493-593 nm. Time series recording spanning 20μm of the islet (Z step: 10 μm, stack of 3 confocal images with a size of 1024 × 1024 pixels) was carried out with a sampling rate of 5 s. A basal recording was acquired for 10 min when the islets were perifused in KRBH buffer with basal glucose concentration of 3 mM. For stimulation, islets were perifused with a KRBH buffer containing 16.7 mM glucose, followed by 3 mM glucose and 16.7 mM glucose plus 60 mM KCl. Data were analyzed using Fiji (NIH)(4) from sum intensity projections. The average Cal-520® AM fluorescence intensity of individual islets was measured in drawn ROIs using Time Series Analyzer plugin (https://imagej.nih.gov/ij/plugins/time-series.html). Changes in cytosolic Ca^2+^ level were calculated and normalized to the average value of Cal-520® AM fluorescence intensity in the baseline (F/F0). Glucose stimulation peak and KCl peak were identified using the integrated measurement of area under the curve in GraphPad Prism v10.3.0 to generate parameters including the onset time, peak time and peak height. The other parameters, including the amplitude, rise time and area of the curve (AOC), were calculated using Microsoft Excel 2016.

### Proximity ligation assay (PLA)

PLA were performed using the Duolink™ In Situ Fluorescence Kit (Sigma-Aldrich, DUO92101) according to the manufacturer’s instructions. Briefly, primary islets were fixed, permeabilized, and incubated with a rabbit anti-villin antibody (Thermo Fisher Scientific, PA5-22072) and a mouse anti-SNAP25 antibody (Synaptic Systems, 111011). After hybridization with PLA probes and rolling-circle amplification, fluorescent PLA signals were visualized by confocal microscopy. Negative controls were processed in parallel using only the anti-villin antibody.

### Statistical methods

Statistical analyses were conducted using GraphPad Prism 10 for macOS (GraphPad Software, San Diego, CA, USA). Two-group comparisons were assessed using Student’s t-test, with unpaired tests used unless otherwise specified in the figure legends. Results are presented as the mean ± standard error (SE) unless otherwise stated. Values of p < 0.05 were considered statistically significant. Bars show standard deviations from at least three independent experiments unless otherwise stated. Histograms were prepared using GraphPad Prism 10 or Microsoft Excel (Microsoft Corp., Redmond, WA). For ipITT, we use coupled test (series of data time dependant) for difference of responses relative to the point t=0

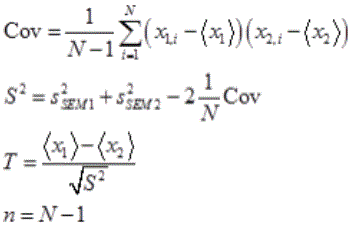

WT –0.00667+-0.134, K/V=-1.13+-0.17

Then we have T = 1.123. Corrected for covariation S = 0.119 and for null-hypothesis that there is no difference is p-value = 0.0007

## Acknowledgments

We thank Katja Pfriem for administrative assistance, Frank Moller and Mark Leaver for assistance in image editing. We also thank Ünal Coskun and Michal Gryzbek from the Center for Membrane Biochemistry and Lipid Research, Faculty of Medicine, TU Dresden, Germany, for their assistance with MST technology.

## This work was supported by

- **German Center for Diabetes Research (DZD e.V.)** Funded by the German Ministry for Education and Research (BMBF).
- **INNODIA and INNODIA HARVEST** This project received funding from the Innovative Medicines Initiative 2 Joint Undertaking under grant agreements No. **115797** (INNODIA) and No. **945268** (INNODIA HARVEST). This Joint Undertaking receives support from the European Union’s Horizon 2020 research and innovation programme, the European Federation of Pharmaceutical Industries and Associations (EFPIA), JDRF (now Breakthrough T1D), and The Leona M. and Harry B. Helmsley Charitable Trust.
- **IRTG 2251: ICMSD** Funding was provided by the Deutsche Forschungsgemeinschaft (DFG, German Research Foundation), Project Number **288034826** (IRTG 2251: ICMSD; MS, PK).
- **INTERCEPT Project** Funded by the European Union’s Horizon Europe research and innovation programme under grant agreement No. **1010954433**.

This research was also partially funded by the Lower Saxony Ministry of Science and Culture (MWK) with funds from the Volkswagen Foundation’s zukunft niedersachsen program [project name: CAIMed – Lower Saxony Center for Artificial Intelligence and Causal Methods in Medicine; grant number: ZN4257]”.

## Required EU Disclaimer

Funded by the European Union. Views and opinions expressed are however those of the author(s) only and do not necessarily reflect those of the European Union. Neither the European Union nor the granting authority can be held responsible for them.

## Figure legends

**Extended Data Figure 1.**
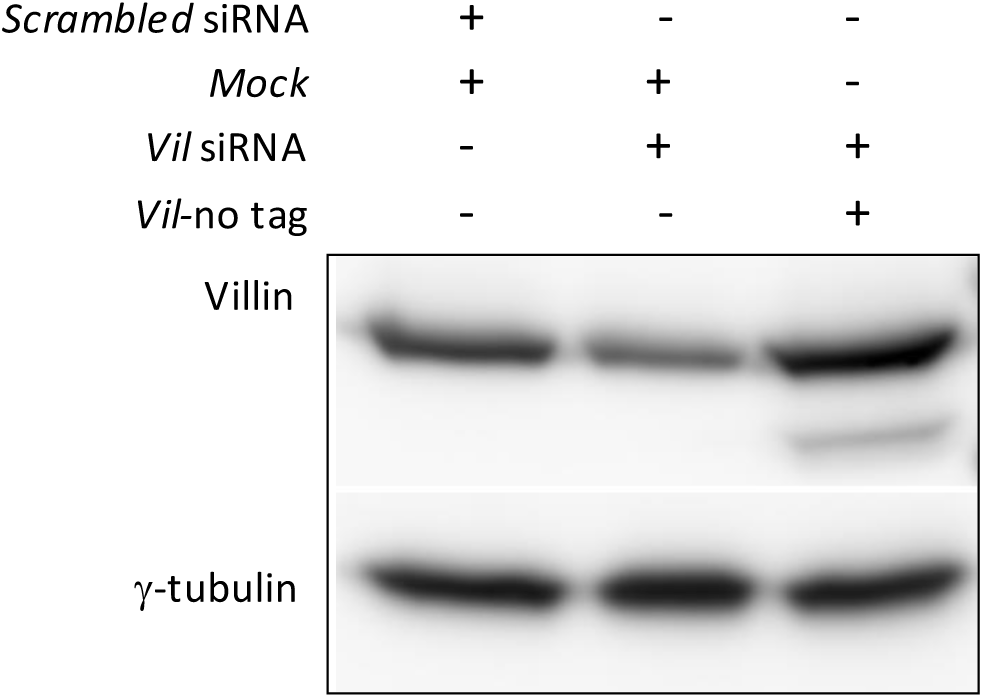
Western blotting for villin and γ-tubulin on extracts from INS-1 cells transfected with *scrambled siRNA + CMV-Mock, Vil* siRNA + *CMV-Mock or Vil* siRNA + *CMV-Vil* in relationship to Fig.1e.

**Extended Data Figure 2.**
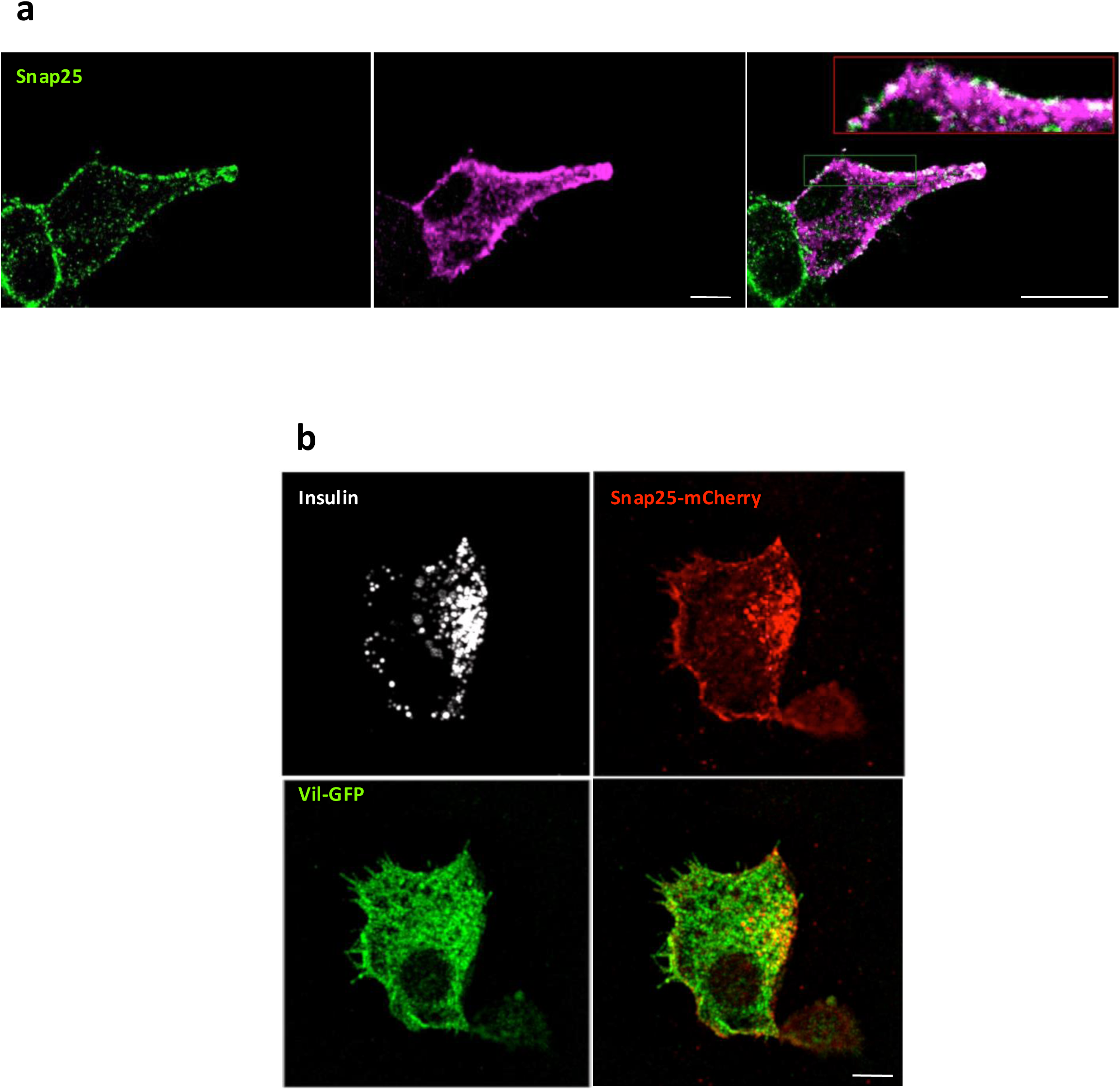
Partial cellular co-localization of villin and SNAP25. (a) confocal images of endogenous villin and SNAP25 in INS-1 cells. The inset is a 2.5 x magnification of the images in (a). (b) High resolution microscopy images of insulin and ectopically expressed SNAP25 and villin in INS-1 cells. Arrows indicate puncta positive for SNAP25 and villin. Scale bars 5 μm in (a) and (b).

**Extended Data Figure 3.**
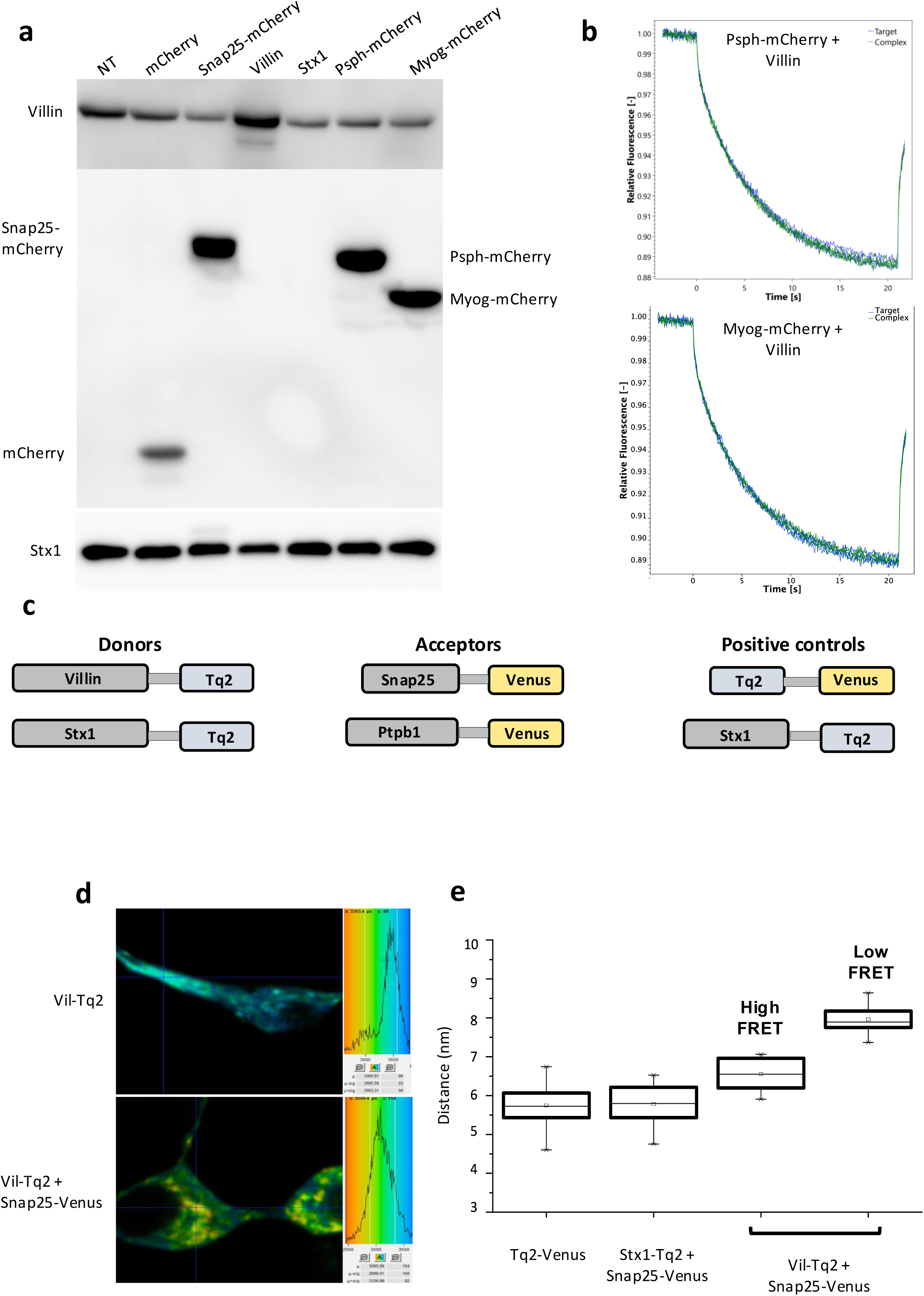
Expression in IN-1 cells of constructs used in MST assay B. (a) Western Blot analysis of INS-1 extracts transfected with indicated constructs and probed with the corresponding antibodies in relationship to experiment Fig.3a. (b) Representative micro scale thermophoresis plots of the fluorescence for mCherry upon mixture of extracts of INS-1 cells transiently transfected with Psph-mCherry or Myog-mCherry with extracts of INS-1 cells transfected with villin as indicated. (c) Schematic representation of the constructs donors, acceptors or controls used in FLIM/FRET. (d) Microscopic images and lifetime plots of donor alone (vil-Tq2) or in presence of an acceptor (Snap25-Venus). (e) Förster distance between SNAP25 and villin in comparison to Snap25-Stx1.

**Extended Data Figure 4.**
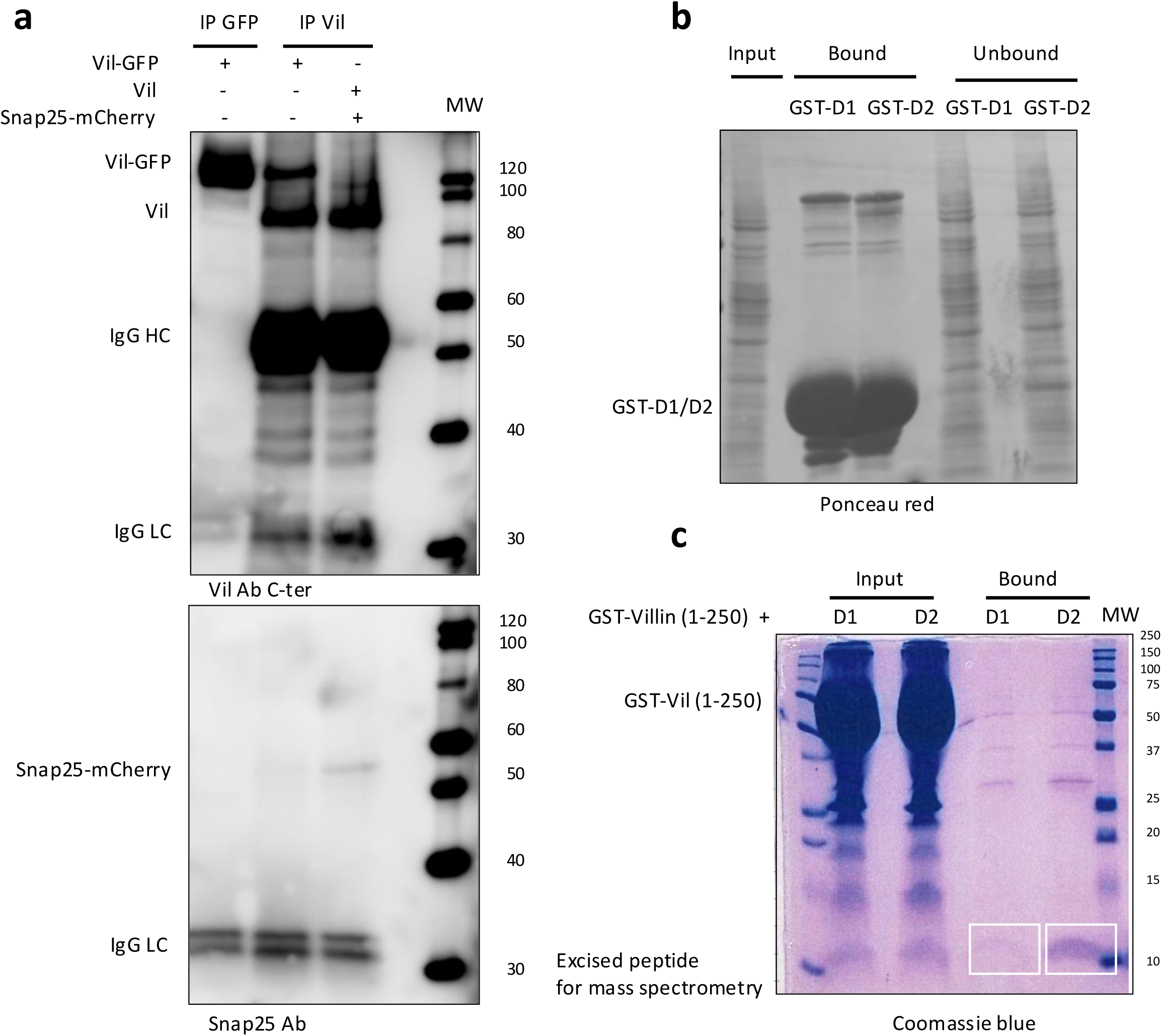
Co-immunoprecipitation of SNAP25 and villin. (a) Co-immunoprecipitation of SNAP25 from cells transfected with non-tagged villin, vil-GFP or SNAP25-mCherry plasmids with control GFP or villin antibodies. The complex was washed and the bound fraction was analyzed by 10% SDS-PAGE and the blots probed with antibodies for villin and SNAP25. (b) Ponceau red staining as control of equal loading of the gel shown in Fig.4c. (c) The N-terminal fragment of villin (amino acids 1-250) expressed as glutathione S-transferase *(GST)* fusion protein was immobilized on glutathione-Sepharose beads. The coiled coil domains D1 (aa:19-81) and D2 (aa:140-202) were expressed in bacteria as GST-fusion proteins and excised with Pre-Scission enzyme and incubated with GST-vil (1-250) in a pull-down assay. The complex was washed and the bound fraction was subjected to 15% SDS-PAGE. The gel stained Coomassie-Blue and the indicated bands were analyzed by Mass Spectrometry.

**Extended Data Figure 5.**
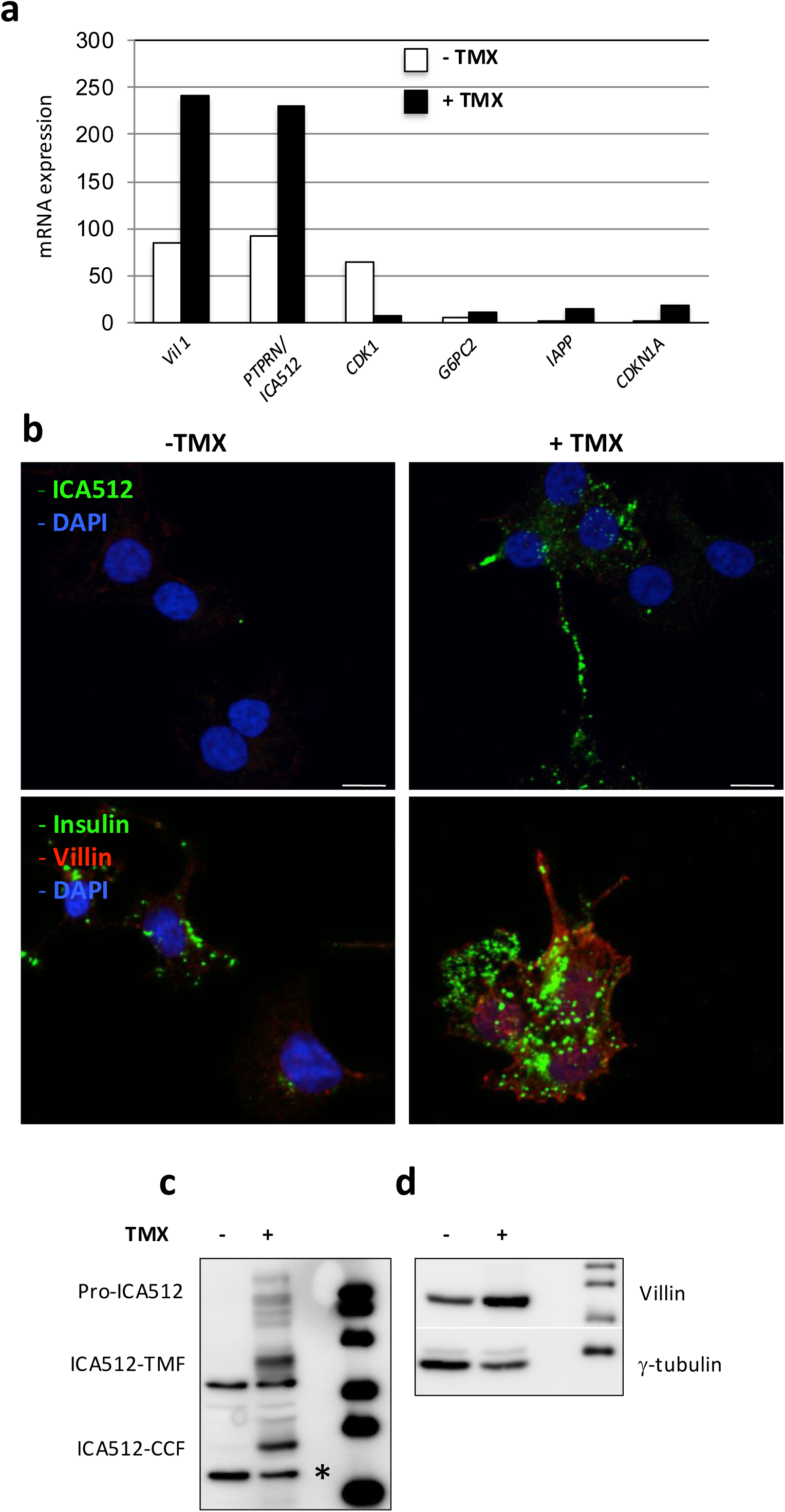
TAM-treatment of EndoC-βH3 correlated with induced expression of ICA512 and villin. (a) Gene profiling analysis of selected β-cell specific genes showed induced expression of ICA512 and villin upon TAM induction of EndoC-βH3. (b) Immunostaining and (b) Western Blot analysis of EndoC-βH3 cells with antibodies for ICA512 (c) and villin (d) prior or after TAM induction. The asterisk corresponds to an unspecific band.

**Extended Data Figure 6.**
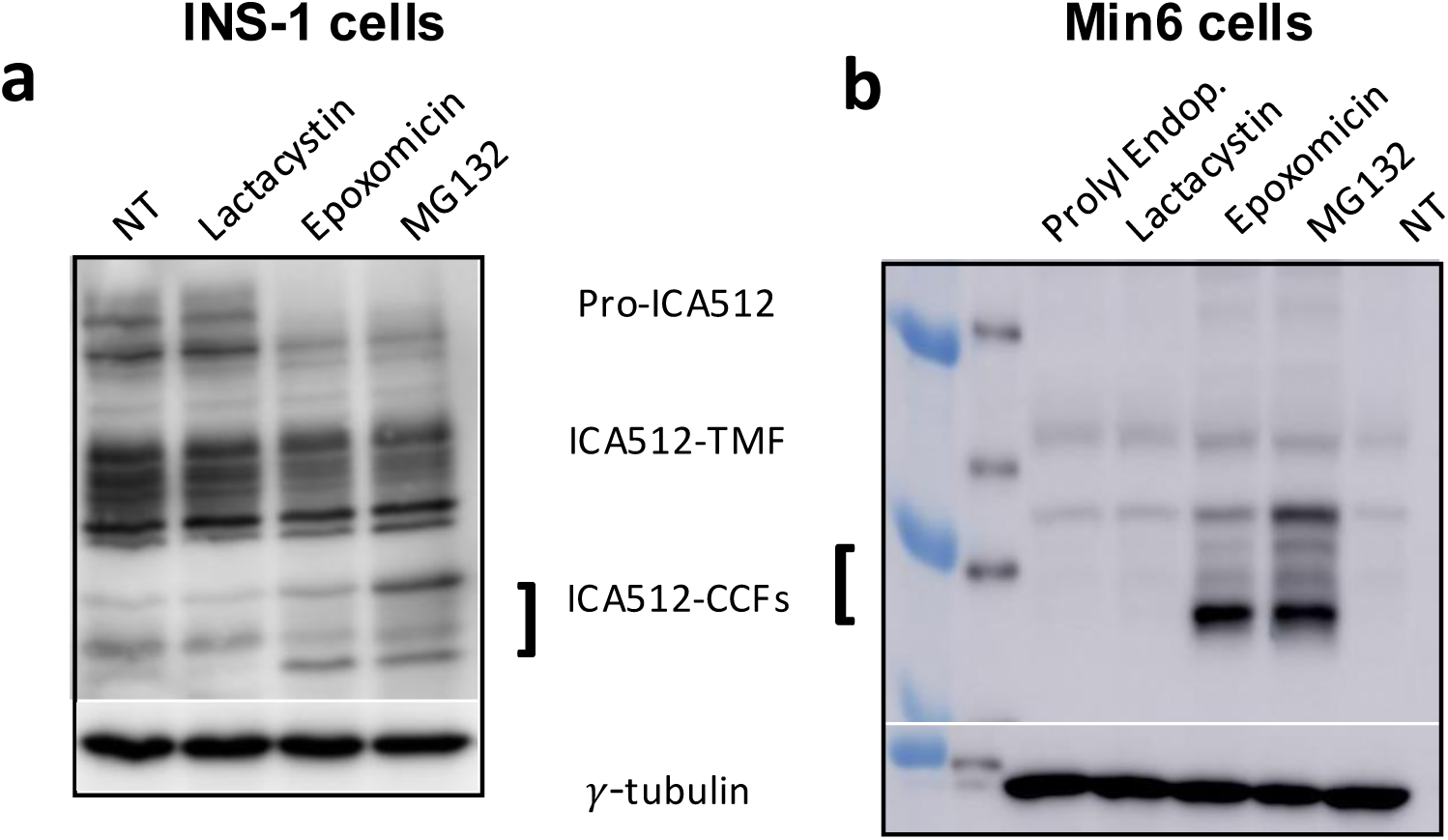
(a-b) Inhibition of ICA512 turn-over and stabilization of ICA512-CCF in INS-1 (a) and MIN6 (b) with the proteasome inhibitors Epoxomicin and MG-132.

**Extended Data Figure 7.**
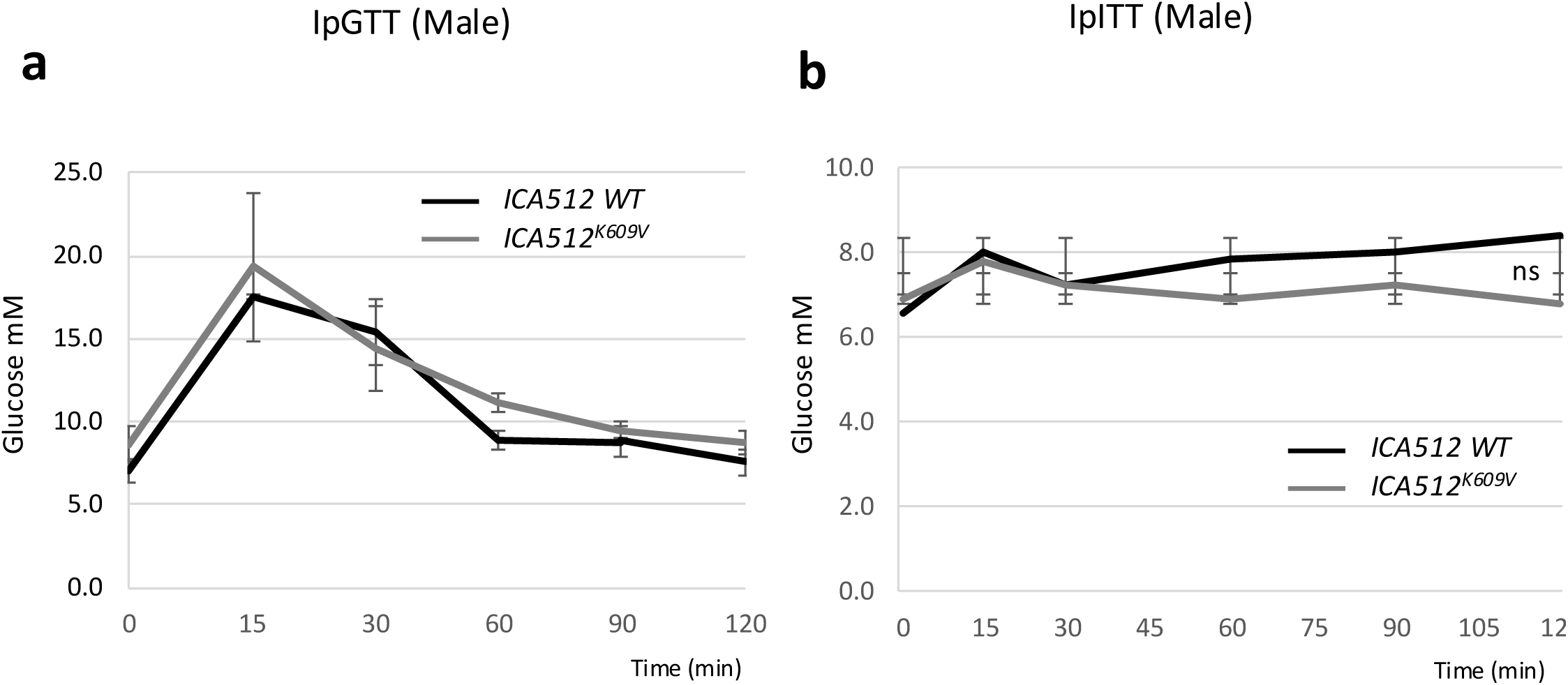
(a) IpGTT on 3 *Ica512^wt^* and 5 *Ica512^K609/V^* male mice. (b) ipITT on 3 *Ica512^wt^* and 7 *Ica512^K609/V^* male mice.

**Extended Data Figure 8.**
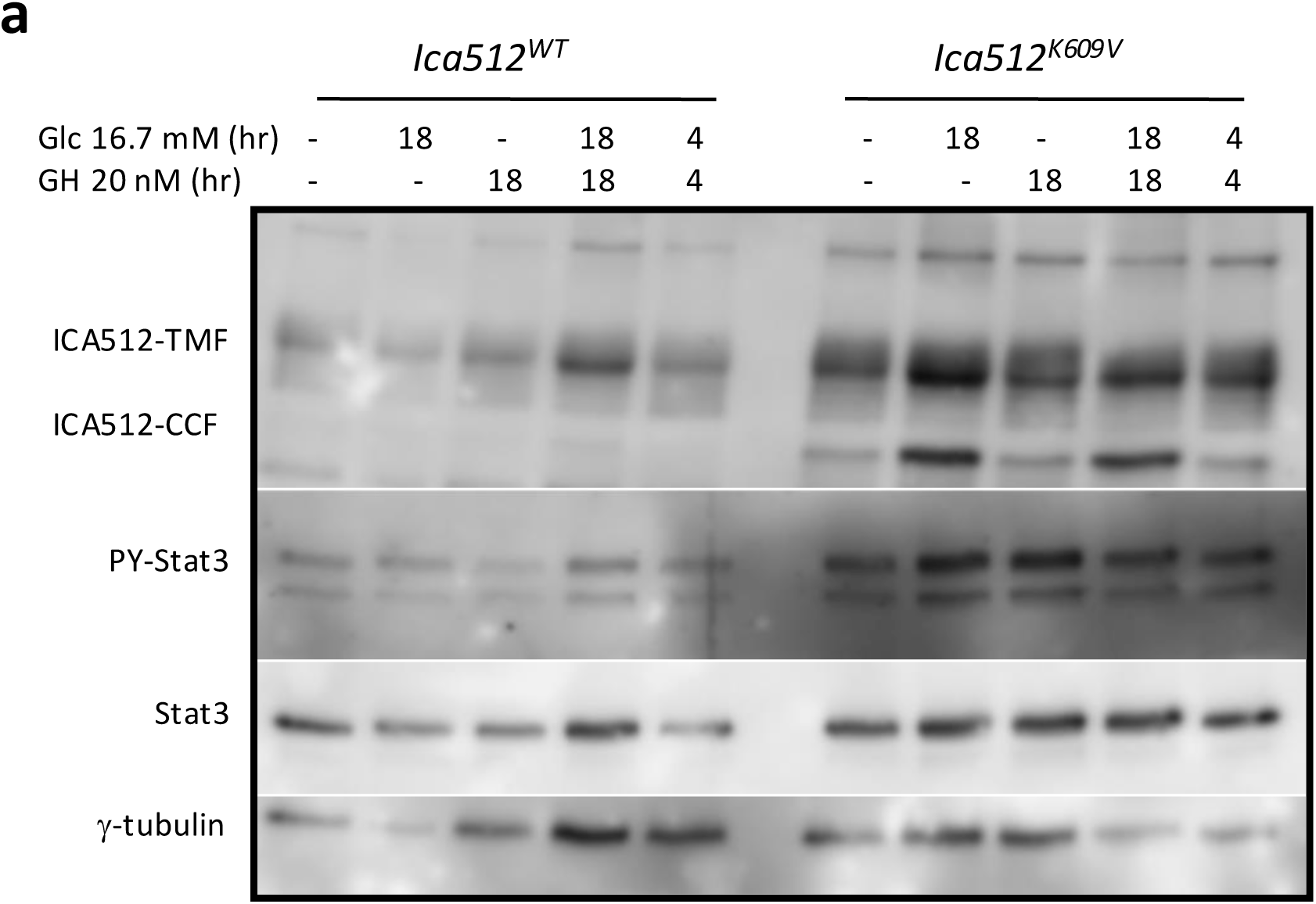
Enhanced STAT3 phosphorylation in *Ica512^K609/V^* mice. Western Blot analysis of islets extracts isolated from *Ica512^wt^* and *Ica512^K609/V^* mice treated as indicated and probed for ICA512, PY-STAT3, STAT3 and γ-tubulin.

**Extended Data Figure 9.**
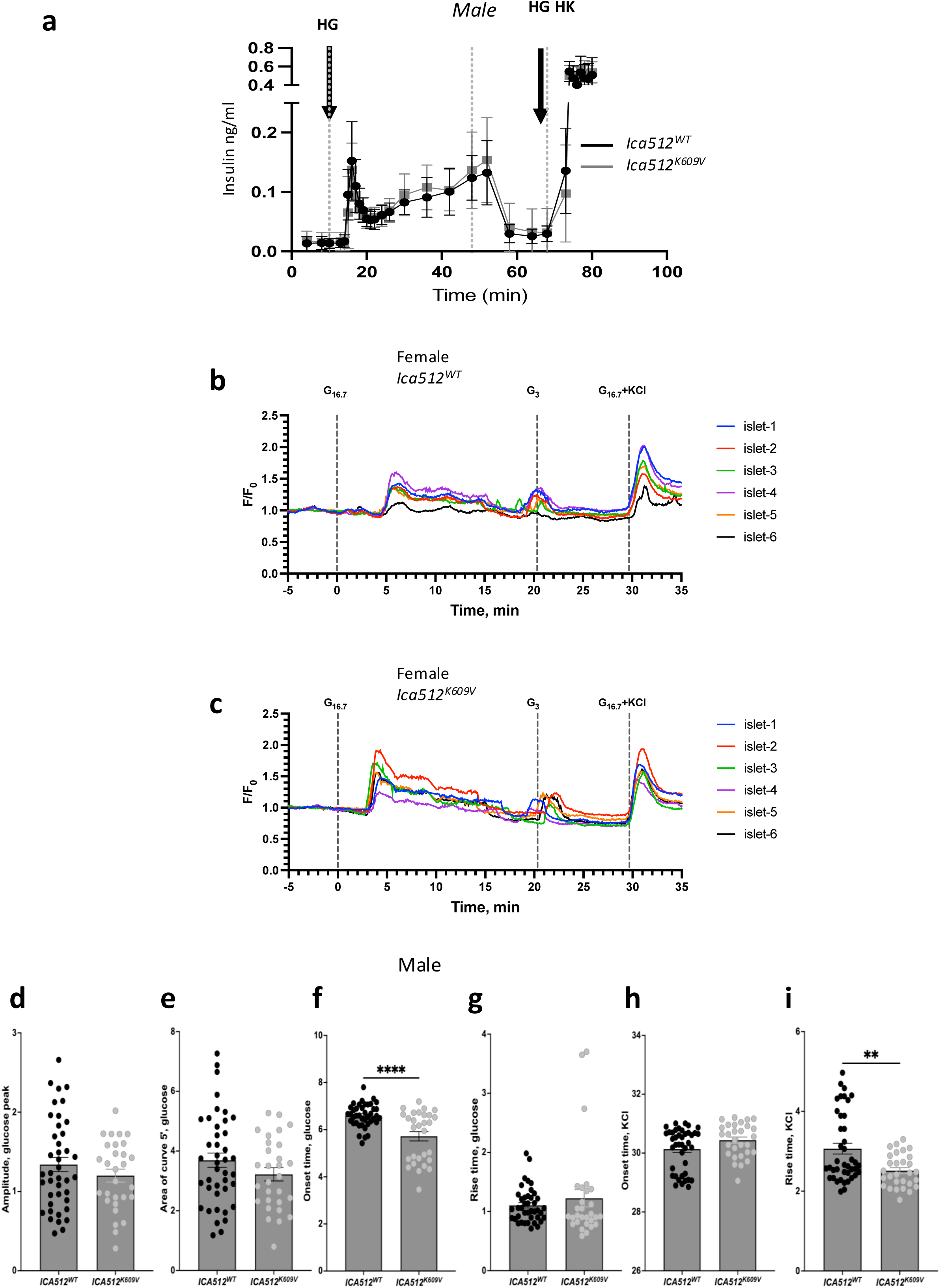
(a) Dynamic GSIS on islets isolated from 3 *Ica512^wt^* and 3 *Ica512^K609/V^* male mice. (b-c) calcium imaging on islets isolated from female mice *Ica512^wt^* and ICA512^K609/V^ treated as in Fig. 8e-i. (d-i) Calcium imaging of glucose– and HKCl-stimulated islets from *Ica512^wt^* and *Ica512^K609/V^* male mice showing reduced onset and rise times respectively after glucose and KCl stimulation. Data in (d-i) are from a group of mice n= 6-7.

**ESM Table 1.** Information on primary and secondary antibodies used for immunoblotting and or for immunostaining.

| **Primary Antibodies** | **Company** | **Cat.#** | **Host** | **Application** | **Dilution** |
| --- | --- | --- | --- | --- | --- |
| Villin-1a | Cell Signaling | 2369 | Rabbit | WB/ICH | 1:1000/1:75 |
| Villin-1b | Thermofisher | PA5-22072 | Rabbit | WB | 1:5000 |
| SNAP25a | Synaptic syst. | 11110 | Mouse | WB | 1:1000 |
| SNAP25 | Synaptic syst. | 111011 | Mouse | WB | 1:1000 |
| ICA512 HM-1 | Home made | HM-1 | Mouse | WB/ICH | 1:500/1:100 |
| Syntaxin1 | Synaptic syst. | 110011 | Mouse | WB/ICH | 1:1000/1:150 |
| Syntaxin4 | Synaptic syst. | 110041 | Mouse | WB | 1:1000 |
| Munc 18 | Cell Signaling | 13414 | Rabbit | WB | 1:1000 |
| mCherry | Clontech | 632543 | Mouse | WB | 1:5000 |
| HA | Abcam | ab9110 | Rabbit | WB | 1:2500 |
| γ-Tubulin | Sigma Aldrich | T6557 | Mouse | WB | 1:5000 |

**ESM Table 2.**
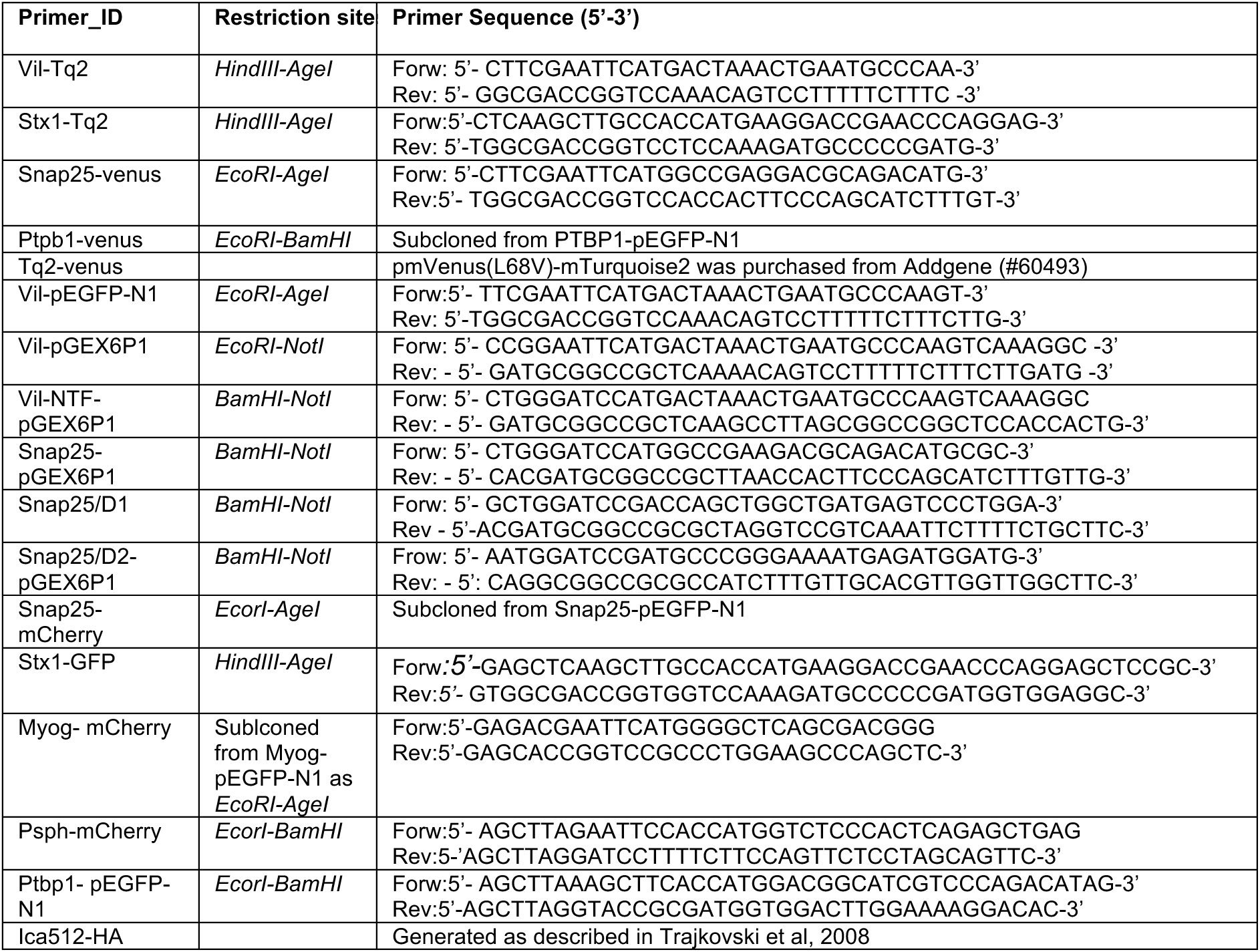
Information on oligonucleotides used for cloning if the constructs used in the manuscript.

